# KAT6A deficiency impairs cognitive functions through suppressing RSPO2/Wnt signaling in hippocampal CA3

**DOI:** 10.1101/2024.03.26.586861

**Authors:** Yongqing Liu, Minghua Fan, Junhua Yang, Ljubica Mihaljević, Kevin Hong Chen, Yingzhi Ye, Shuying Sun, Zhaozhu Qiu

## Abstract

Intellectual disability (ID) affects ∼2% of the general population and is often genetic in origin. ID-associated genes are enriched for epigenetic factors, including those encoding the largest family of histone lysine acetyltransferases (KAT5-KAT8). Among them is *KAT6A*, whose *de novo* heterozygous mutations cause KAT6A Syndrome (or Arboleda-Tham Syndrome), with ID as a common clinical feature. However, the underlying molecular mechanisms remain elusive. Here, we show that haploinsufficiency of *Kat6a* impairs learning and memory in mice, and specific deletion of *Kat6a* in excitatory neurons recapitulates the hippocampus-dependent memory deficits. Unexpectedly, KAT6A deficiency results in impaired synaptic structure and plasticity in hippocampal CA3, but not in CA1 region. Combining single-nucleus RNA-sequencing and chromatin analysis, we identify a CA3-enriched gene *Rspo2*, encoding a Wnt activator R-spondin 2, as a key transcriptional target of KAT6A. Moreover, deletion of *Rspo2* in excitatory neurons phenocopies the loss of *Kat6a*, resulting in defective Wnt/β-catenin signaling and synaptic plasticity in CA3, and abnormal cognitive behaviors in mice. Importantly, restoring RSPO2 expression in CA3 pyramidal neurons rescues the deficits in Wnt signaling and learning-associated behaviors in *Kat6a* mutant mice. Collectively, our results demonstrate that KAT6A plays a critical role in regulating synaptic plasticity and memory formation through RSPO2-mediated Wnt signaling in hippocampal CA3, shedding new light on the fundamental mechanisms of ID and providing potential therapeutic targets for the treatment of KAT6A Syndrome and related neurodevelopmental diseases.

## INTRODUCTION

The development of central nervous system and the learning-induced rewiring of neural circuits are tightly modulated through the dynamic regulation of gene expression (*1, 2*). Acetylation of histones is a key mechanism regulating chromatin organization and gene expression (*3*). While histone deacetyltransferases (HDACs) have been the focus of intense research, the role of histone acetyltransferases (HATs) in the brain is under-explored. The five-member MYST proteins represent the largest family of HATs in humans, whose gene mutations often cause neurodevelopmental diseases characterized with intellectual disability (ID) (*4–8*).

One of MYST family member KAT6A (also known as MOZ and MYST3) was first identified as an oncogene rearranged in acute myeloid leukemia (*9*). It is composed of a double PHD finger that binds to acetylated histone tails, the HAT domain, and a long unstructured C-terminal region. KAT6A plays an important role in tumor growth, body segment patterning, and hematopoietic stem cell maintenance (*10–12*). Heterozygous nonsense (protein-truncating, most likely loss-of-function) mutations of *KAT6A* also cause a rare genetic disorder KAT6A Syndrome (or Arboleda-Tham Syndrome), with ∼400 patients reported so far around the world (*13*). With the increased application of whole-exome sequencing, the number of diagnosed individuals will likely grow much larger. KAT6A Syndrome is characterized by several common clinical traits including ID, speech and language deficits, and global developmental delay, along with less penetrant phenotypes such as microcephaly, neonatal hypotonia, and autism spectrum disorder (*13*). Most of the pathogenic variants for KAT6A Syndrome are protein-truncating mutations that occur throughout the gene. However, patients with mutations in last two exons (exon 16-17) have more severe ID and other phenotypes, compared to those with early-truncating mutations (*13*). It’s possible that these late-truncating mutations are not subjected to nonsense-mediated mRNA decay. The resulting truncated KAT6A proteins may possess dominant negative effects. Nevertheless, the impact of *KAT6A* loss-of-function in the nervous system and the underlying mechanism leading to the brain disease remain elusive.

Here, we uncover a KAT6A/RSPO2/Wnt signaling cascade that is required for learning and memory by regulating synaptic structure and plasticity in hippocampal CA3 pyramidal neurons, and provide new insights to our understanding of the pathogenesis of KAT6A Syndrome.

## RESULTS

### *Kat6a* haploinsufficiency results in cognitive deficits in mice

To elucidate the mechanism behind KAT6A Syndrome and assess the effect of KAT6A deletion in the brain, we first generated *Kat6a*-floxed mice using the efficient additions with ssDNA inserts-CRISPR (*Easi*-CRISPR) method (*14*) (**Fig. 1A** and **S1A**). During the insertion of a floxed cassette, we also obtained *Kat6a* knockout allele because of deletion of the target exon (**Fig. 1A** and **S1B-C**). *Kat6a* homozygous knockout mice were embryonic lethal (*12*), hence we tested haploinsufficiency in *Kat6a*^+/-^ mice with exon 4 deletion that results in a premature stop codon. Therefore, it probably mimics patients with early-truncating mutations. The mRNA level of *Kat6a* in hippocampus of *Kat6a*^+/-^ mice was ∼50% of that in wild-type (WT) littermates (**Fig. 1B**). Both male and female *Kat6a*^+/-^ mice displayed smaller body weight than WT littermates starting at weaning age and throughout adulthood (**Fig. 1C** and **S1D-E**), consistent with growth delays of patients with KAT6A Syndrome.

**Fig. 1.**
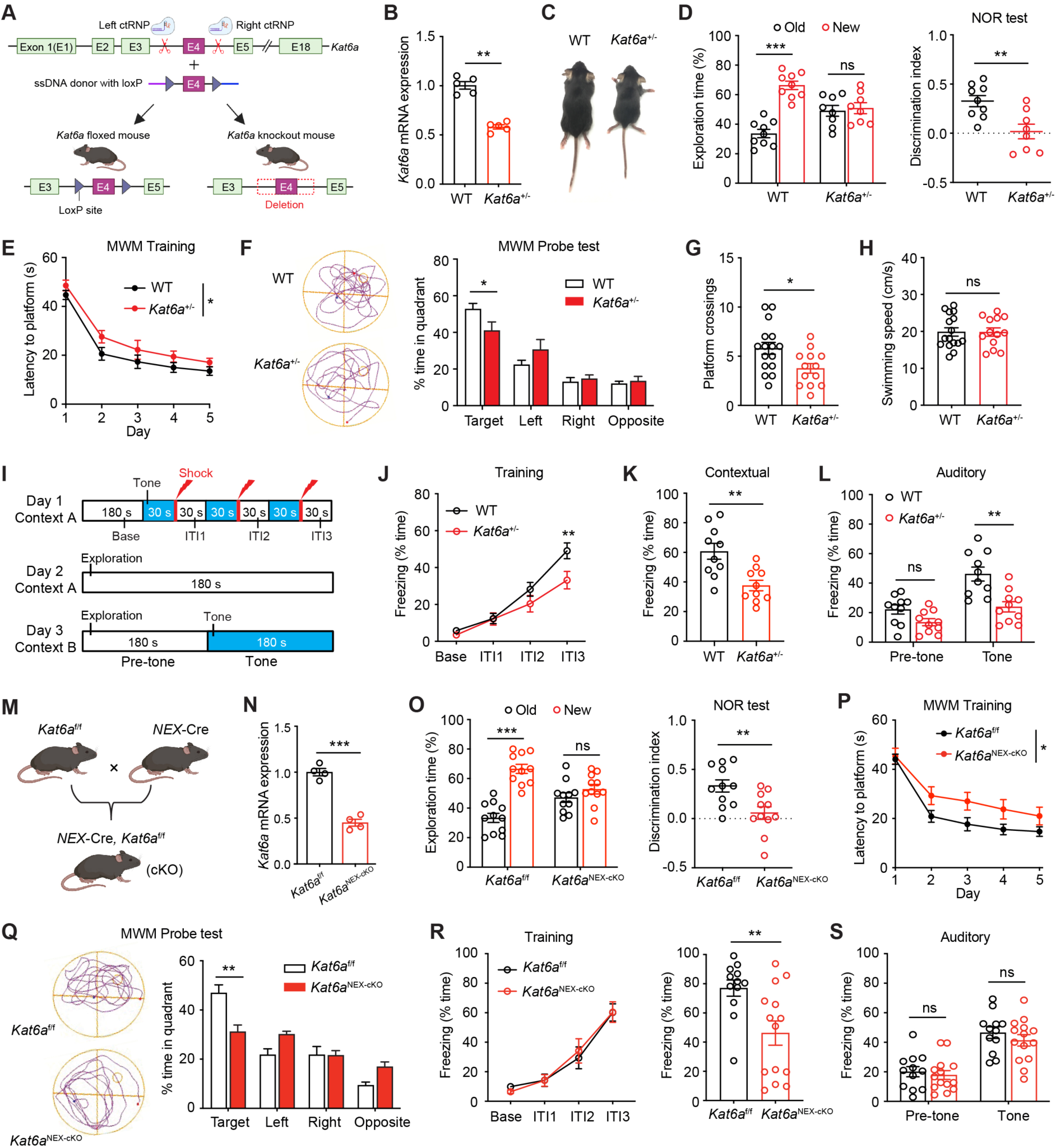
KAT6A deficiency impairs learning and memory in mice. (**A**) Schematic diagram. (**B**) qRT-PCR analysis of total hippocampal mRNA from WT and *Kat6a*^+/-^ mice. n = 5 mice per group. (**C**) Representative images of body size of 4-week-old WT and *Kat6a*^+/-^ mice. (**D**) The NOR test. n = 8-9 mice per group. (**E**) Time spent before reaching the hidden platform during training days in the MWM test. Two-way ANOVA test. F (1, 27) = 5.645, *p = 0.025. n = 13-16 mice per group. (**F**) Representative traces and time spent in different quadrants during the probe test. (**G** and **H)** Entries into platform zone and swimming speed in the MWM test. Unpaired Student’s t test. (**I**) Schematic diagram of fear conditioning test. ITI, inter-trial interval. (**J**) Learning curves during the training sessions in fear conditioning test. Two-way ANOVA test. n = 10 mice per group. (**K**) Contextual test. Unpaired Student’s t test. (**L**) Auditory test. Two-way ANOVA test. (**M**) *Kat6a*^f/f^ mice crossed with *NEX*-cre mice to obtain the *Kat6a*^NEX-cKO^ mice. (**N**) qRT-PCR analysis of total hippocampal mRNA from *Kat6a*^f/f^ and *Kat6a*^NEX-cKO^ mice. n = 4 mice per group. (**O**) The NOR test. n = 11 mice per group. (**P**) Time spent before reaching the hidden platform during training days in the MWM test. Two-way ANOVA test. F (1, 20) = 6.713, *p = 0.017. (**Q**) The probe test of the MWM test. n = 11 mice per group. (**R**) Left: Learning curves during the training sessions in fear conditioning test. Two-way ANOVA test. F (1, 24) = 0.004, p = 0.952. Right: Contextual test. n = 12 *Kat6a*^f/f^ and n = 14 *Kat6a*^NEX-cKO^ mice. Unpaired Student’s t test. (**S**) Auditory test. *p < 0.05, **p < 0.01, ***p < 0.001. ns, nonsignificant. Data are presented as mean ± SEM.

We then performed a battery of behavioral assays to compare the performance of *Kat6a*^+/-^ and their WT littermates in adulthood. In the open field test, *Kat6a*^+/-^ mice showed more time spent at the center of the chamber compared to control mice, whereas both groups showed comparable total distance traveled (**Fig. S1F**), indicating that the mutant mice have normal locomotor activity but with less anxiety levels. The reduced anxiety was further confirmed by the elevated plus maze test (**Fig. S1G**). In addition, we used the three-chamber assay to examine the sociability of WT and *Kat6a*^+/-^ mice. When compared with WT mice, the majority of *Kat6a*^+/-^ mice exhibited normal social ability (**Fig. S1H-I**), consistent with the notion that many parents describe their children with KAT6A mutations as happy and sociable despite ID and developmental delays.

Considering that ID is a prominent clinical feature of KAT6A Syndrome, we further investigated whether KAT6A loss affects cognitive functions in mice. In the novel object recognition (NOR) test, which evaluates the preference of mice to explore a new object over a familiar object, WT littermates displayed a significant preference for the novel object (**Fig. 1D**). However, *Kat6a*^+/-^ mice showed no such preference (**Fig. 1D**), indicating a deficit in recognition memory.

To test hippocampus-dependent spatial learning and memory, we used the Morris water maze (MWM) assay to test an animal’s ability to use spatial cues to locate a hidden platform in a tank of water. During the training sessions, *Kat6a*^+/-^ mice spent a longer time navigating to the platform than control mice (**Fig. 1E**), indicating that their spatial learning is impaired. In the probe test, *Kat6a*^+/-^ mice spent less time in the target quadrant (**Fig. 1F-G**), whereas their swimming activity was normal (**Fig. 1H**), suggesting spatial memory is impaired in the mutant mice.

To further confirm the importance of KAT6A in learning and memory, we performed a classic fear conditioning procedure, in which we paired an auditory cue with a mild aversive foot shock (**Fig. 1I**). Contextual fear memory is sensitive to hippocampal defects, whereas auditory fear memory depends on amygdala function (*15*). During the training sessions, *Kat6a*^+/-^ mice exhibited decreased foot shock-evoked freezing behavior compared with WT controls (**Fig. 1J**), suggesting that their learning ability is impaired. At twenty-four hours after conditioning, mice were placed back into the training context and their freezing time was measured. Compared with controls, *Kat6a*^+/-^ mice displayed significantly decreased context-evoked freezing behaviors (**Fig. 1K**). Likewise, decreased tone-evoked freezing behaviors were also observed in the mutants (**Fig. 1L**). Together, these data demonstrate that *Kat6a* haploinsufficiency results in memory deficits in mice.

### Excitatory neuron-specific deletion of KAT6A is sufficient to cause memory deficits

In the central nervous system, KAT6A is ubiquitously expressed across different cell types. To explore which cell type in the brain is susceptible to the loss of KAT6A function in the context of cognitive defects, we first crossed *Kat6a* floxed mice (*Kat6a*^f/f^) with *NEX*-cre mice that express Cre recombinase in pyramidal neurons in the neocortex and hippocampus (*Kat6a*^NEX-cKO^) (**Fig. 1M**). In the *Kat6a*^NEX-cKO^ mice, the level of *Kat6a* mRNA was significantly reduced in hippocampus (**Fig. 1N**). Notably, compared to control littermates, adult *Kat6a*^NEX-cKO^ mice appeared grossly normal, with normal cortical morphology (**Fig. S2A-C**), hippocampal cell density (**Fig. S2D**), body weight (**Fig. S2E**), and adult hippocampal neurogenesis (**Fig. S2F**). However, similar to *Kat6a*^+/-^, *Kat6a*^NEX-cKO^ mice also displayed reduced anxiety levels (**Fig. S2G-H**). Moreover, *Kat6a*^NEX-cKO^ mice exhibited memory deficits in the NOR test and the MWM test (**Fig. 1O-Q**), suggesting that mice with KAT6A deletion in excitatory neurons recapitulate the memory deficits in *Kat6a*^+/-^ mice. In the fear conditioning test, compared with control mice, both male (**Fig. 1R**) and female (**Fig. S2I**) *Kat6a*^NEX-cKO^ mice displayed significantly decreased context-evoked freezing behavior, whereas there was no difference in baseline freezing in a new context and no difference in elevated freezing due to the tone presentation (**Fig. 1S** and **S2I**), indicating that memory deficits of *Kat6a*^NEX-cKO^ mice are specific to hippocampus-dependent contextual fear conditioning.

To bypass any potential developmental effects and further validate that excitatory neuronal loss of KAT6A leads to cognitive deficits, we carried out a second cross with the *Camk2a*-cre mice (**Fig. S3A**), where Cre expression starts at 2∼3 weeks after birth and is restricted to forebrain excitatory neurons in hippocampus and cortex. Similar to *Kat6a*^NEX-cKO^, *Kat6a*^Camk2a-cKO^ mice also displayed memory defects in the learning-related behaviors (**Fig. S3**), while showing normal locomotion, anxiety level, and sociability (**Fig. S3**). In contrast, ablation of KAT6A in astrocytes by *Gfap*-cre (**Fig. S4A-E**) or in inhibitory neurons by *Gad2*-cre (**Fig. S4F**) did not induce cognitive dysfunction. Together, these results demonstrate that loss of KAT6A specifically in excitatory neurons is sufficient to cause memory deficits in mice.

### KAT6A deficiency impairs hippocampal CA3 synaptic plasticity

To investigate whether deletion of KAT6A influences neuronal function in mouse hippocampus, a pivotal region implicated in learning and memory, we performed whole-cell patch-clamp recordings on acute hippocampal slices to first examine the basal synaptic transmission of CA1 pyramidal neurons (**Fig. 2A**). Surprisingly, miniature excitatory postsynaptic currents (mEPSCs) of CA1 pyramidal neurons were comparable in frequency and amplitude between *Kat6a*^NEX-cKO^ mice and control littermates (**Fig. 2B**). In addition, both paired-pulse ratios (**Fig. 2C**) and long-term potentiation (LTP) (**Fig. 2D**) in dorsal CA1 region were normal in *Kat6a*^NEX-cKO^ mice, suggesting that KAT6A deficiency has no obvious effect on CA1 neuronal function.

**Fig. 2.**
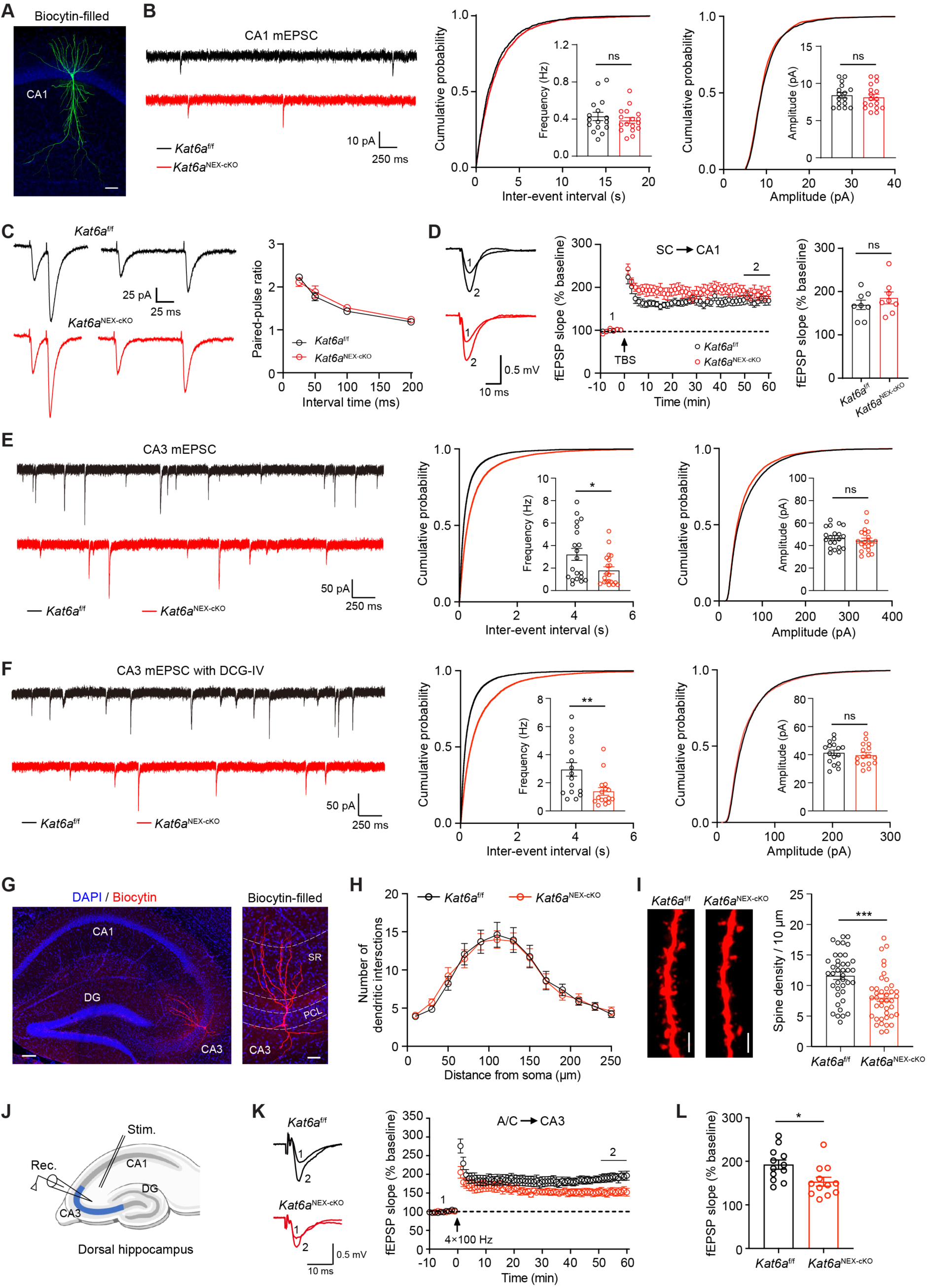
Loss of KAT6A results in hippocampal CA3 synaptic plasticity defects. (**A**) Representative biocytin-filled CA1 pyramidal neurons from *Kat6a*^f/f^ mice. Scale bar, 50 µm. (**B**) Representative traces (left) and quantification of frequency (middle) and amplitude (right) of mEPSC in dorsal CA1 pyramidal neurons. n = 16-17 cells from 4 mice per group. Unpaired Student’s t test. (**C**) PPRs with different interstimulus intervals. n = 16 cells from 3 mice per group. Two-way ANOVA test. (**D**) TBS-induced LTP at Schaffer collateral (SC) to CA1 synapses. Arrow indicates LTP induction. n = 8 slices from 4 mice per group. Unpaired Student’s t test. (**E**) mEPSC in dorsal CA3 pyramidal neurons. n = 20 cells from 4 mice per group. Mann-Whitney test. (**F**) mEPSC in the presence of DCG-IV in dorsal CA3 pyramidal neurons. n = 16 cells from 4 mice per group. Mann-Whitney test. (**G**) Left: Representative biocytin-filled CA3 pyramidal neurons from *Kat6a*^f/f^ mice. Scale bar, 100 µm. Right: Different layers in CA3 region. Scale bar, 40 µm. SR, stratum radiatum. PCL, pyramidal cell layer. (**H**) Quantification of CA3 pyramidal neuron dendritic intersections revealed by Sholl analysis. n = 13 cells from 4 mice per group. (**I**) Left: Representative images of dendritic segments in the stratum radiatum dendrites of CA3 pyramidal neurons. Scale bar, 2 µm. Right: Quantification of dendritic spine numbers. n = 40 dendrites from 4 mice per group. Unpaired Student’s t test. (**J**) Illustration of LTP at CA3 recurrent synapses. (**K**) 4×HFS-induced LTP in the presence of DCG-IV at A/C to CA3 synapses. (**L**) The averaged fEPSP slopes during 50-60 min after the stimulation. n = 12 slices from 4 mice per group. Unpaired Student’s t test. *p < 0.05, **p < 0.01, ***p < 0.001. ns, nonsignificant. Data are presented as mean ± SEM.

Given that the hippocampal CA3 region is also implicated in memory function (*16–18*), we next measured mEPSC in CA3 pyramidal neurons. Interestingly, mEPSC frequency was substantially decreased in *Kat6a*^NEX-cKO^ mice compared to controls, whereas amplitude was not changed (**Fig. 2E**), suggesting that KAT6A deficiency causes defects in CA3 glutamatergic synaptic transmission. CA3 pyramidal neurons receive three main types of glutamatergic inputs: recurrent CA3 collaterals, mossy fiber (MF) inputs from DG granule cells, and perforant path inputs from entorhinal cortex (*18*). To further examine which input contributes to this defect, we recorded mEPSCs while blocking MF inputs and partial perforant path inputs using the group II metabotropic glutamate receptor agonist DCG-IV (*19, 20*). In the presence of DCG-IV, a significant decrease in mEPSC frequency was still observed in CA3 pyramidal neurons of *Kat6a*^NEX-cKO^ mice (**Fig. 2F**), indicating that CA3 recurrent synaptic transmission is mainly impaired in the absence of KAT6A.

To explore the underlying mechanisms responsible for the defects in synaptic transmission upon deletion of *Kat6a*, we filled CA3 pyramidal neurons with biocytin and imaged their dendritic branches (**Fig. 2G**). While the dendrite arbors were not changed in *Kat6a*^NEX-cKO^ mice by Sholl analysis (**Fig. 2H**), we observed a significant decrease in spine density at CA3 stratum radiatum, a main region receiving CA3 recurrent inputs (**Fig. 2I**). Furthermore, in the presence of DCG-IV, LTP induced by high-frequency stimulation at associational and commissural (A/C) fibers to CA3 synapses of the mutant mice was partially but significantly reduced (**Fig. 2J-L**). Collectively, the above data suggest that deletion of KAT6A in excitatory neurons impairs hippocampal CA3 synaptic plasticity, thus contributing to the learning and memory deficits.

### snRNA-seq reveals transcriptional targets of KAT6A in hippocampal CA3

To decipher the molecular mechanisms of *Kat6a* haploinsufficiency-induced memory defects, RNA sequencing (RNA-seq) was performed with total RNA extracted from hippocampus of *Kat6a*^+/-^ mice and their WT littermates. A cutoff of *p* value < 0.05 and |log_2_ FC| > 0.2 was applied to identify differentially expressed genes (**Fig. 3A**). Because KAT6A mainly acts as a transcriptional coactivator, we focused on gene downregulation, as these genes are more likely to be its direct targets. A total of 185 genes were downregulated in hippocampus of *Kat6a*^+/-^ mice (**Fig. 3B**), which were classified into various signaling pathways using Gene Ontology (GO) enrichment analysis with a cutoff of *q* value < 0.05. Many of the downregulated genes are involved in the Wnt signaling pathway and excitatory synapse function (**Fig. 3C** and **Table S1**), supporting the idea that KAT6A plays an important role in developmental processes and regulation of synaptic plasticity.

**Fig. 3.**
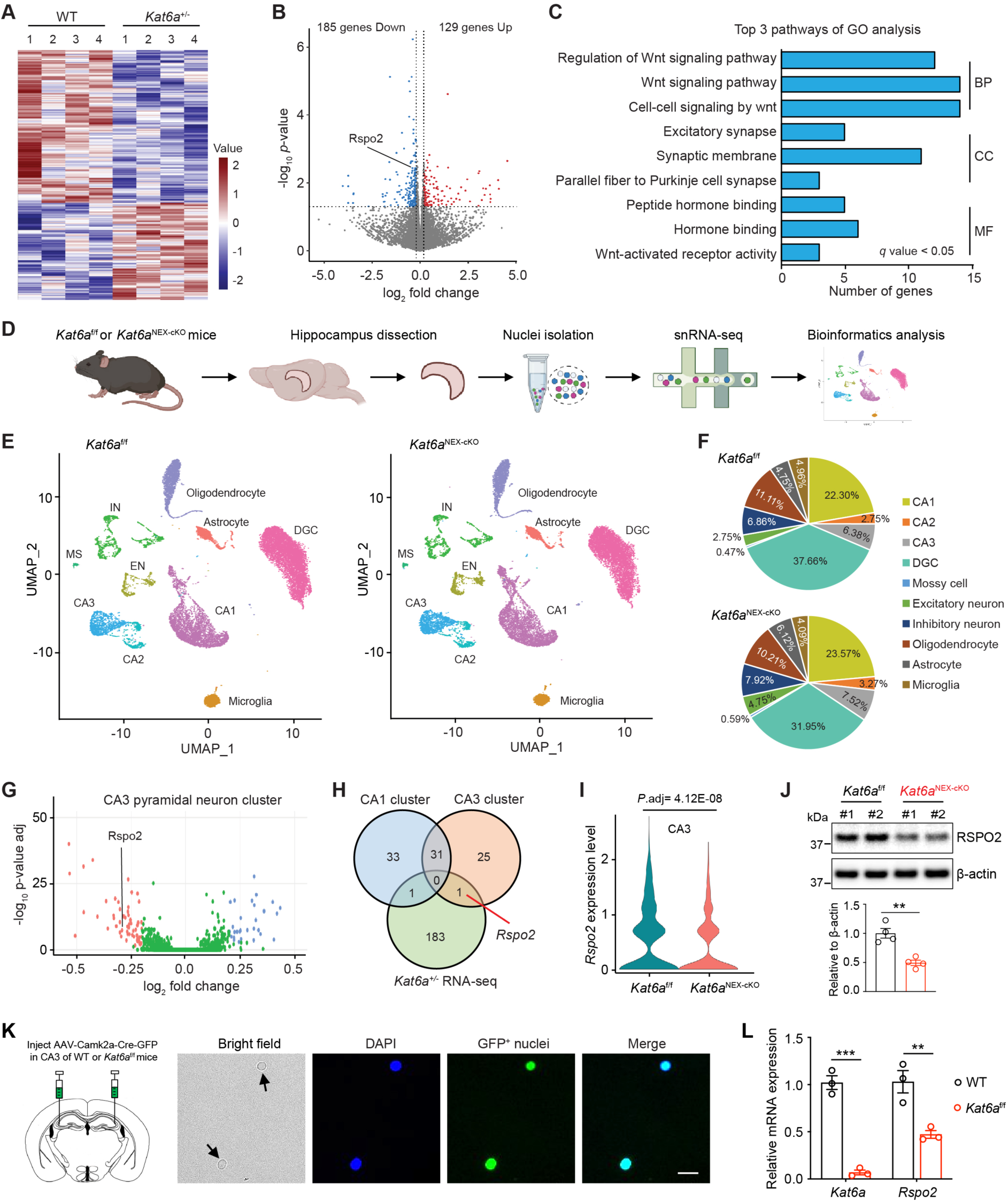
Identification of *Rspo2* as a key target gene of KAT6A in hippocampal CA3. (**A**) Hippocampus tissues were obtained from adult male WT and *Kat6a*^+/-^ mice, followed by RNA extraction and deep sequencing. Heatmap showing up- or downregulated genes indicated by red or blue, respectively. (**B**) Volcano plot showing differentially expressed genes (WT versus *Kat6a*^+/-^). (**C**) Classification of the downregulated genes in *Kat6a*^+/-^ mice with GO analysis. BP, biological process. CC, cellular component. MF, molecular function. (**D**) Flow chart of snRNA-seq using hippocampi from adult male *Kat6a*^f/f^ mice and *Kat6a*^NEX-cKO^ mice. (**E**) UMAP plot of cell clusters from snRNA-seq. (**F**) Percentile plots of the major hippocampal cell types from snRNA-seq in *Kat6a*^f/f^ mice and *Kat6a*^NEX-cKO^ mice. (**G**) Volcano plot showing differentially expressed genes in CA3 pyramidal neuron cluster (*Kat6a*^f/f^ versus *Kat6a*^NEX-cKO^). (**H**) Venn diagram of overlapping downregulated genes among CA1 pyramidal neuron cluster, CA3 pyramidal neuron cluster and hippocampal RNA-seq of *Kat6a*^+/-^ mice. (**I**) Violin plot showing expression levels of *Rspo2* in CA3 pyramidal cells from *Kat6a*^f/f^ mice and *Kat6a*^NEX-cKO^ mice. Unpaired Student’s t test. **(J**) Western blotting showing deletion of KAT6A decreased RSPO2 expression in hippocampal CA3. Quantification was done by normalizing to β-actin. n = 4 mice, unpaired Student’s t test. (**K**) Scheme (left) of AAV-Camk2a-Cre-GFP virus injection into the mouse hippocampal CA3. Representative images (right) of the GFP^+^ nuclei from mouse hippocampus after AAV injection. Scale bar, 5 µm. (**L**) qRT-PCR analysis of GFP^+^ enriched mRNA from WT and *Kat6a*^f/f^ mice for *Kat6a* and *Rspo2*. Two-way ANOVA test. **p < 0.01, ***p < 0.001. ns, nonsignificant. Data are presented as mean ± SEM.

As cell-type-specific changes in the transcriptome can be difficult to detect using bulk RNA-seq in the brain tissues that consist of complex mixtures of cells, we next performed single-nucleus RNA sequencing (snRNA-seq) to more precisely unravel which genes are changed upon *Kat6a* deletion in excitatory neurons. Hippocampal nuclei from *Kat6a*^NEX-cKO^ and *Kat6a*^f/f^ mice were isolated and subsequently sorted using the 10x Genomics platform (**Fig. 3D**). Transcripts were sequenced from 18,421 and 17,806 nuclei from hippocampi of *Kat6a*^NEX-cKO^ and control mice, respectively. The Uniform Manifold Approximation and Projection (UMAP) analysis identified all of the major cell types in hippocampus based on their unique gene-expression signatures (**Fig. 3E** and **S5**). No apparent difference in cell type distribution was observed in both groups (**Fig. 3F**), in line with the normal adult hippocampal neurogenesis in *Kat6a*^NEX-cKO^ mice (**Fig. S2F**). Given our finding that KAT6A deficiency results in cognitive deficits and CA3 synaptic dysfunction, we paid specific attention to the differentially expressed genes (*q* value < 0.05, and |log_2_ FC| > 0.2) in the CA3 pyramidal neuron cluster (**Fig. 3G**). Interestingly, based on the analysis of the transcripts from CA3 cluster of *Kat6a*^NEX-cKO^ mice and hippocampal RNA-seq of *Kat6a*^+/-^ mice, only one common gene (R-spondin 2, *Rspo2*) was significantly downregulated in both groups, whereas this gene was not changed in CA1 pyramidal neurons and other cell types (**Fig. 3H-I** and **Fig. S6A-B**). This suggests that *Rspo2* is a promising candidate gene involved in KAT6A-mediated cognitive functions, although other gene expression changes could also play a role in this process (**Fig. S6C**).

RSPO2, belonging to R-spondin family secreted proteins (RSPO1-4), is an activator of Wnt/β-catenin signaling and is involved in development and cancer (*21–23*). It is highly expressed in mouse hippocampus (**Fig. S6D**). Consistent with the sequencing data, *Rspo2* expression level was decreased in hippocampi from both *Kat6a*^NEX-cKO^ (**Fig. 3J** and **Fig. S6E**) and *Kat6a*^+/-^ mice (**Fig. S6F**). To further confirm that *Rspo2* downregulation occurs in CA3 pyramidal neurons, we stereotactically injected AAV viruses expressing Cre-GFP into hippocampal CA3 of *Kat6a*^f/f^ mice. At four weeks after virus injection, hippocampal nuclei were isolated and subsequently sorted to enrich the GFP^+^ nuclei (**Fig. 3K**). Downregulation of *Rspo2* in CA3 excitatory neurons upon KAT6A deletion was validated by qRT-PCR assays (**Fig. 3L**).

To test if KAT6A directly regulates *Rspo2* transcription, we next performed chromatin immunoprecipitation (ChIP) experiments. Due to the lack of a ChIP-grade antibody against KAT6A, we generated and validated a *Kat6a* knock-in mouse strain with a 3×HA tag fused to its C-terminus (KAT6A-HA) (**Fig. 4A-B**). *Kat6a*^HA/HA^ mice are viable and healthy, suggesting that the KAT6A-HA fusion protein is functional. Using ChIP-qPCR with an HA antibody, strong enrichment of KAT6A was observed at the promoter region of *Rspo2* in hippocampus (**Fig. 4C**). Significantly, knockout of *Kat6a* in excitatory neurons decreased enrichment of histone H3 lysine 23 acetylation (H3K23ac), but not H3K9ac or H3K14ac (**Fig. 4D**), in line with the previous studies that KAT6A and its paralog KAT6B were mainly responsible for H3K23ac (*24, 25*). Together, these results suggest that KAT6A regulates *Rspo2* transcription by modulating H3K23 acetylation on its promoter.

**Fig. 4.**
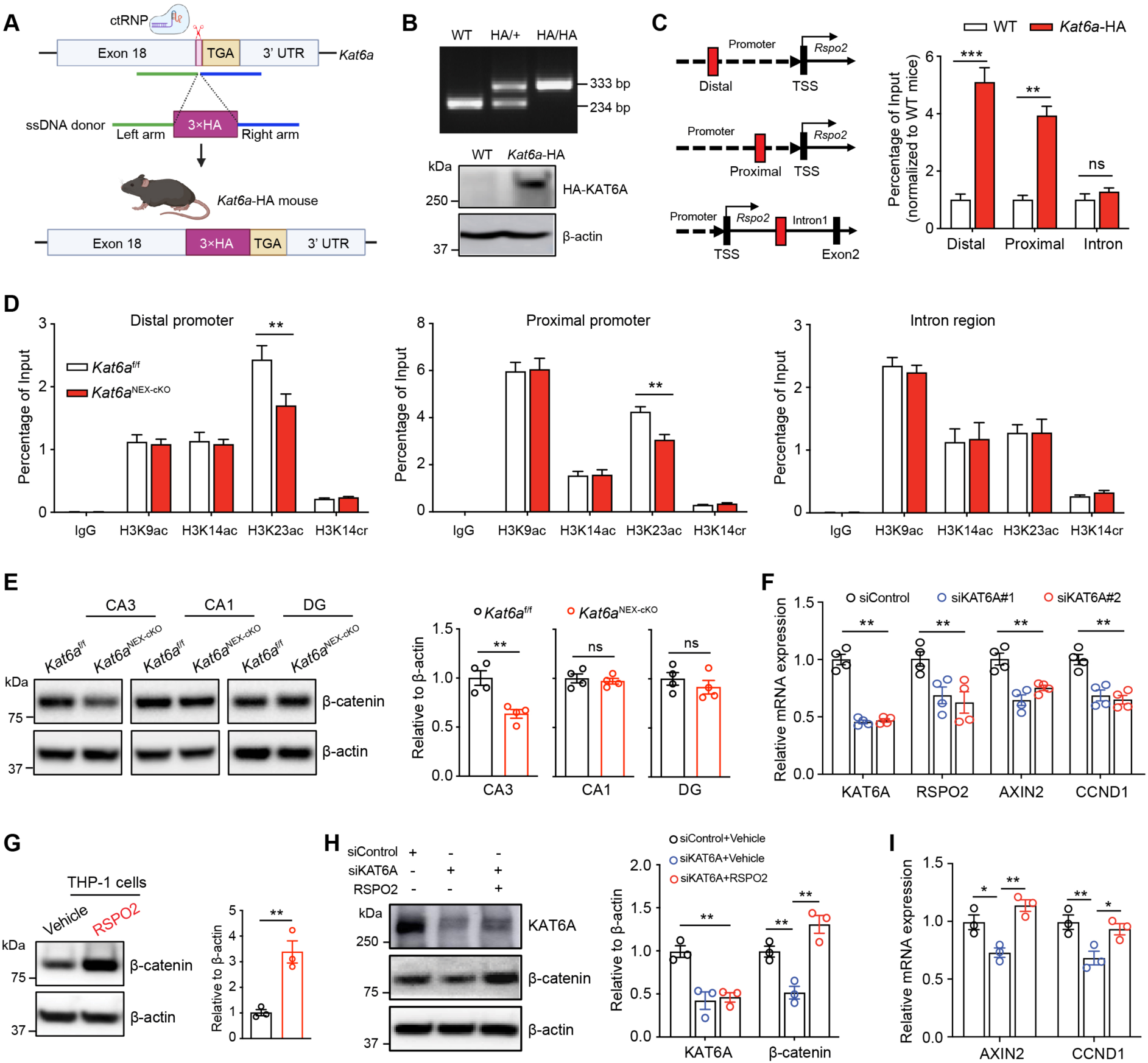
KAT6A deficiency impairs RSPO2-mediated Wnt signaling. (**A**) Scheme of *Kat6a*-HA mice generation. (**B**) Validation of *Kat6a*-HA mice by mouse tail genotyping and western blotting analysis using mouse hippocampal tissues. (**C**) ChIP-qPCR experiments were performed using HA antibody in hippocampus of WT and *Kat6a*-HA mice. Profiles of KAT6A-HA along the mouse *Rspo2* promoter in ∼1500 bp (distal) and ∼300 bp (proximal) upstream of the transcription start site (TSS). n = 3, two-way ANOVA test. (**D**) Deletion of KAT6A resulted in decreased regional enrichment of H3K23ac on the *Rspo2* promoter. Lysates from hippocampal tissues of *Kat6a*^f/f^ mice and *Kat6a*^NEX-cKO^ mice were collected, and ChIP-qPCR assays were performed using the indicated antibodies. n = 3, two-way ANOVA test. (**E**) Western blotting in hippocampal different regions. Quantification was done by normalizing to β-actin. n = 4, unpaired Student’s t test. (**F**) qRT-PCR analysis showing knockdown of *KAT6A* decreased *RSPO2* and Wnt target genes *AXIN2* and *CCND1* expression in human THP-1 cells. mRNA levels were normalized to *GAPDH*. n = 4, two-way ANOVA test. (**G**) Western blotting demonstrating RSPO2 recombinant protein treatment increased active β-catenin expression in human THP-1 cells. Quantification was done by normalizing the level of β-catenin to that of β-actin. n = 3, unpaired Student’s t test. (**H**) Western blotting showing that RSPO2 incubation could counteract the decrease of active β-catenin induced by knockdown of *KAT6A*. Quantification was done by normalizing the level of β-catenin to that of β-actin. n = 3, two-way ANOVA test. (**I**) qRT-PCR analysis showing that RSPO2 incubation reverses the decrease of Wnt target genes *AXIN2* and *CCND1* induced by knockdown of *KAT6A*. mRNA levels were normalized to *GAPDH*. n = 3, two-way ANOVA test. *p < 0.05, **p < 0.01, ***p < 0.001. ns, nonsignificant. Data are presented as mean ± SEM.

### KAT6A regulates RSPO2-mediated Wnt signaling

RSPO2 is a secreted protein and functions as a Wnt agonist to potentiate Wnt/β-catenin signaling by binding to the leucine-rich repeat-containing G-protein-coupled receptors 4/5/6 (LGR4/5/6) (*21, 26*). This prompted us to examine whether KAT6A modulates Wnt signaling in hippocampal CA3 by influencing *Rspo2* expression. Indeed, upon ablation of KAT6A, we observed a significant decrease of active β-catenin in protein lysates from CA3, but not in CA1 or dentate gyrus (DG) (**Fig. 4E**). To further dissect this new genetic pathway, we employed human monocyte THP-1 cell lines as a model, which displays high endogenous expression of RSPO2 (*27*). Consistently, knockdown of KAT6A markedly reduced RSPO2 expression (**Fig. 4F**). It also downregulated AXIN2 and CCND1, two classic Wnt target genes, indicating a decrease in Wnt signaling (**Fig. 4F**). Conversely, treating THP-1 cells with recombinant RSPO2 protein led to an increase in active β-catenin (**Fig. 4G**), consistent with a previous study (*27*). While KAT6A downregulation resulted in a significant decrease in active β-catenin and Wnt target gene (AXIN2 and CCND1) expression, RSPO2 recombinant protein treatment completely reversed these effects (**Fig. 4H-I**). These findings demonstrate that RSPO2 is necessary for KAT6A-mediated regulation of Wnt signaling.

### RSPO2 is essential for hippocampal CA3 synaptic functions

To further examine the expression and function of RSPO2 in hippocampus, we first performed *in situ* hybridization and observed that *Rspo2* was preferentially expressed in the CA3 pyramidal neurons with low expression levels in ventral CA1 neurons (**Fig. 5A** and **Fig. S7A-D**), consistent with our snRNA-seq results (**Fig. 5B**). This distinct expression pattern of *Rspo2* in hippocampus suggests a unique role in CA3 pyramidal neurons. To directly test it, we crossed *Rspo2*^f/f^ mice with *NEX*-cre line to generate excitatory neuron-specific *Rspo2* knockout mice (*Rspo2*^NEX-cKO^). Depletion of *Rspo2* mRNA in hippocampus was verified by qRT-PCR assays (**Fig. 5C**). *Rspo2*^NEX-cKO^ mice were viable and appeared grossly normal (**Fig. S7E**). The active β-catenin level in hippocampal CA3 of *Rspo2*^NEX-cKO^ mice was markedly reduced, whereas the non-canonical Wnt/JNK signaling was normal (**Fig. 5D**), suggesting that RSPO2 is a *bona fide* enhancer of Wnt/β-catenin signaling in hippocampus.

**Fig. 5.**
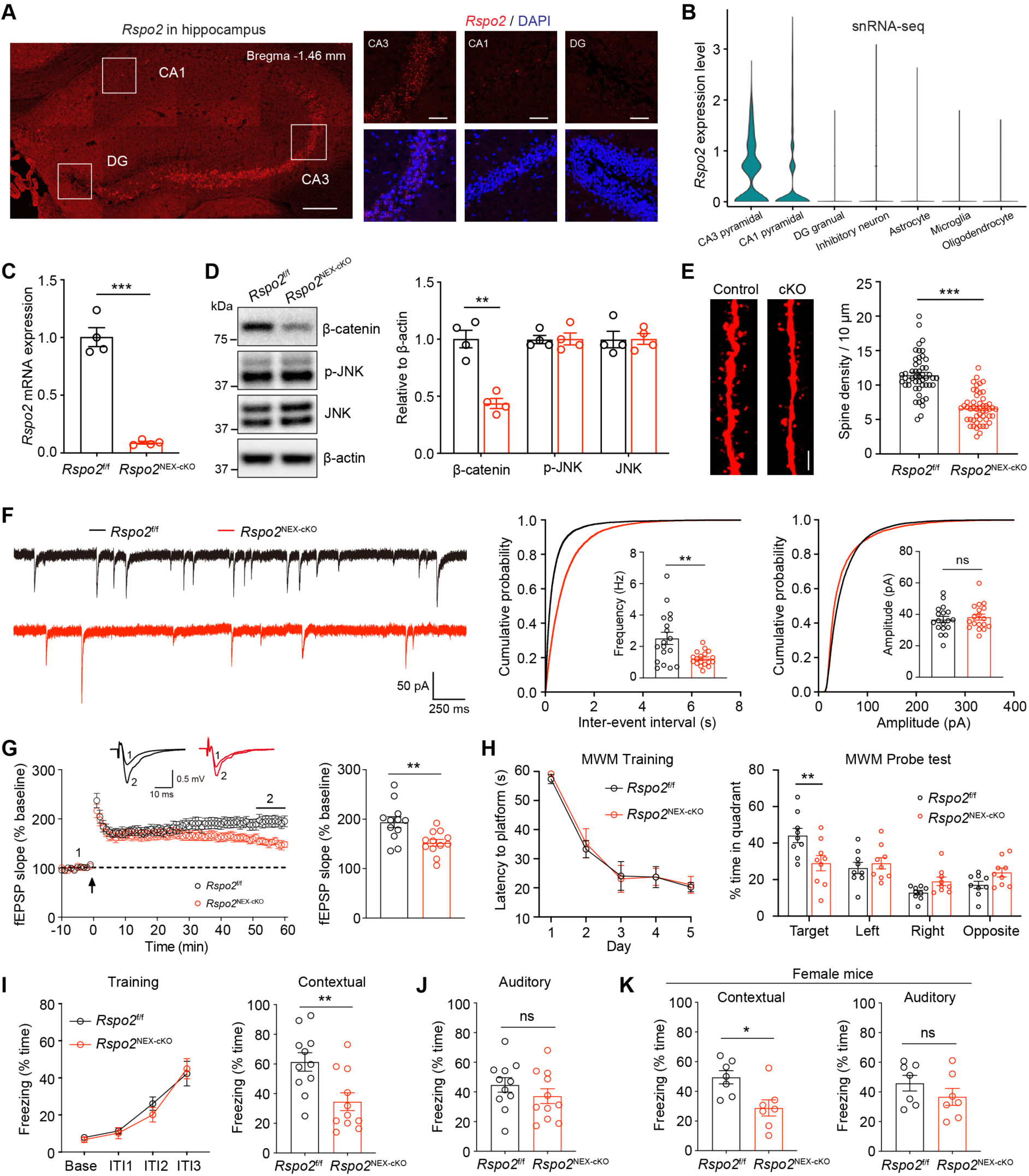
RSPO2 is essential for hippocampal CA3 synaptic functions. (**A**) Representative images of *Rspo2* RNAscope *in situ* hybridization in mouse dorsal hippocampus. Scale bar, 200 µm (left), 40 µm (right). (**B**) Expression levels of *Rspo2* in hippocampal different cell types from the snRNA-seq of *Kat6a*^f/f^ mice. (**C**) qRT-PCR analysis of total hippocampal mRNA showing that *Rspo2* was deleted in *Rspo2*^NEX-cKO^ mice. mRNA levels were normalized to *Gapdh*. n = 4, unpaired Student’s t test. (**D**) Western blotting showing that deletion of RSPO2 decreased active β-catenin expression in hippocampal CA3. Quantification was done by normalizing to β-actin. n = 4, two-way ANOVA test. (**E**) Left: Representative images of dendritic segments in the stratum radiatum dendrites of CA3 pyramidal neurons. Scale bar, 2 µm. Right: Quantification of dendritic spine numbers. n = 50 dendrites from 4 mice per group. Unpaired Student’s t test. (**F**) mEPSC in dorsal CA3 pyramidal neurons. n = 18 cells from 4 mice per group. Unpaired Student’s t test. (**G**) 4×HFS-induced LTP in the presence of DCG-IV at A/C to CA3 synapses. n = 12 slices from 4 mice per group. Unpaired Student’s t test. (**H**) The MWM test. n = 9 mice per group. Two-way ANOVA test. (**I**) Left: Learning curves of male *Rspo2*^f/f^ and *Rspo2*^NEX-cKO^ mice during the training sessions in fear conditioning test. Two-way ANOVA test. F (1, 20) = 0.131, p = 0.721. Right: Contextual test in the training context. n = 11 mice per group. Unpaired Student’s t test. (**J**) Auditory test in a novel context. Unpaired Student’s t test. (**K**) Female *Rspo2*^f/f^ mice and *Rspo2*^NEX-cKO^ mice were used in fear conditioning test. n = 7 mice per group. Unpaired Student’s t test. *p < 0.05, **p < 0.01, ***p < 0.001. ns, nonsignificant. Data are presented as mean ± SEM.

We next examined the effect of RSPO2 deficiency on the morphology and function of CA3 pyramidal neurons. Remarkably, just like *Kat6a^NEX-cKO^* mice, spine density at stratum radiatum of CA3 pyramidal neurons was decreased in *Rspo2*^NEX-cKO^ mice compared to controls (**Fig. 5E**). Correspondingly, we observed a decreased mEPSC frequency, whereas the amplitude of mEPSC was normal (**Fig. 5F**). In addition, LTP induced by high-frequency stimulation at A/C fibers to CA3 synapses of *Rspo2*^NEX-cKO^ mice was also reduced (**Fig. 5G**). These data suggest that similar to KAT6A, the CA3-enriched RSPO2 is critical for hippocampal CA3 synaptic structure and plasticity.

To explore whether the defective synaptic transmission and plasticity lead to any abnormal animal behaviors, we performed the MWM test to evaluate hippocampus-dependent spatial learning and memory in *Rspo2*^NEX-cKO^ mice. While they showed a normal learning ability during the training sessions, in the probe test, the mutant mice spent less time in the target quadrant compared to control mice (**Fig. 5H**). Moreover, the mutant mice displayed decreased context-evoked freezing in the fear conditioning test, with no difference in the tone-evoked freezing behaviors (**Fig. 5I-K**), indicating that the memory deficits of *Rspo2*^NEX-cKO^ mice are hippocampus-dependent. In addition, loss of RSPO2 did not alter the motor abilities or anxiety levels (**Fig. S7F-G**). Together, these data suggest that RSPO2 deficiency phenocopies the loss of KAT6A, impairing CA3 synaptic functions and disrupting memory formation in hippocampus.

### RSPO2 is required for KAT6A-mediated cognitive functions

Given that *Rspo2* is a main transcriptional target of KAT6A in hippocampal CA3 and that RSPO2 depletion impairs memory formation, we hypothesized that RSPO2 is required for KAT6A-dependent synaptic functions. To test this hypothesis, we delivered AAVs expressing either RSPO2-FLAG (AAV-RSPO2) or a control fluorescent protein GFP (AAV-GFP) to the hippocampal CA3 neurons of *Kat6a*^NEX-cKO^ or control mice by bilateral stereotactic injection (**Fig. 6A**). At five weeks after virus injection, the expression efficiency of RSPO2 protein in CA3 was confirmed by immunostaining (**Fig. 6B**). Overexpressing the Wnt agonist RSPO2 in the CA3 pyramidal neurons counteracted KAT6A deletion-induced β-catenin decrease in *Kat6a*^NEX-cKO^ mice (**Fig. 6C**). Moreover, it enhanced mEPSC frequency and increased the dendritic spine density of these mutant neurons to WT control levels (**Fig. 6D-E** and **S8A**), indicating that KAT6A-RSPO2 axis is important for hippocampal CA3 synaptic functions.

**Fig. 6.**
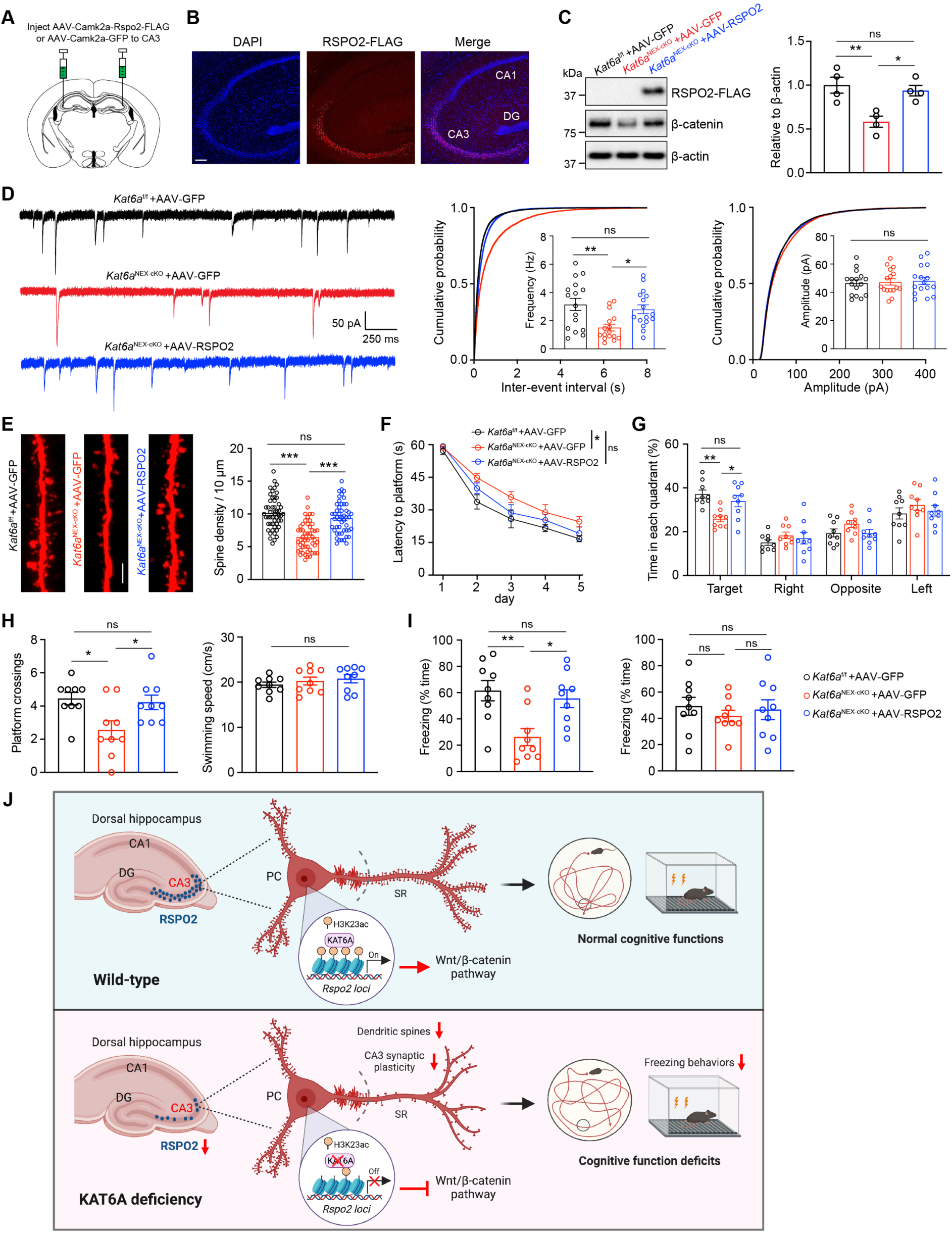
RSPO2 is required for KAT6A-mediated cognitive functions. (**A**) Scheme of AAV virus injection. (**B**) Immunofluorescence assays showing the expression of AAV-Camk2a-Rspo2-FLAG viruses in 5 weeks after infecting the mouse CA3 area. Scale bar, 100 µm. (**C**) Western blotting showing that infection of AAV-RSPO2 could counteract the decrease of active β-catenin induced by deletion of KAT6A. Quantification was done by normalizing to β-actin. n = 4, one-way ANOVA test. (**D**) Representative traces (left) and quantification of frequency (middle) and amplitude (right) of mEPSC in dorsal CA3 pyramidal neurons of *Kat6a*^f/f^ mice injected with AAV-GFP, *Kat6a*^NEX-cKO^ mice injected with AAV-GFP, and *Kat6a*^NEX-cKO^ mice injected with AAV-RSPO2. n = 16 cells from 4 mice per group. One-way ANOVA test. (**E**) Left: Representative z-stack images of dendritic segments in the stratum radiatum dendrites of CA3 pyramidal neurons. Scale bar, 2 µm. Right: Quantification of dendritic spine numbers. n = 50 dendrites from 4 mice per group. One-way ANOVA test. (**F**) Time spent before reaching the hidden platform during training days in the MWM test. Two-way ANOVA test. n = 9 mice for each group. (**G**) Time spent in different quadrants during the probe test of the MWM test. Two-way ANOVA test. (**H**) Entries into platform zone (left) and swimming speed (right) in the MWM test. One-way ANOVA test. (**I**) Contextual (left) and auditory (right) tests in fear conditioning assays. n = 9 mice for each group. One-way ANOVA test. (**J**) Summary diagram showing that the KAT6A-RSPO2 axis is required for memory formation by regulating synaptic structure and plasticity in hippocampal CA3 pyramidal cells (PCs). *p < 0.05, **p < 0.01, ***p < 0.001. ns, nonsignificant. Data are presented as mean ± SEM.

We further performed a proof-of-concept study to examine whether RSPO2 overexpression ameliorates KAT6A deficiency-induced cognitive defects. Remarkably, injection of AAV-RSPO2 in the CA3 of *Kat6a*^NEX-cKO^ mice almost completely rescued the learning and memory deficits in the MWM test (**Fig. 6F-H** and **S8B**). Furthermore, in the fear conditioning test, while no difference in the tone-evoked freezing was observed in these mice, overexpression of RSPO2 in hippocampal CA3 neurons largely alleviated the context-evoked freezing defects caused by KAT6A deletion (**Fig. 6I** and **S8C**). Together, both functional and behavioral data demonstrate that RSPO2 is indispensable for KAT6A-mediated synaptic and cognitive functions and suggest that memory deficits can be reversed in the *Kat6a* mutant mice by elevating canonical Wnt signaling pathway.

## DISCUSSION

Our work identifies histone acetyltransferase KAT6A as a critical regulator of hippocampal CA3 synaptic function by modulating RSPO2/Wnt signaling. It reveals the impairment of KAT6A/RSPO2/Wnt signaling pathway as a key molecular mechanism underlying cognitive dysfunction caused by KAT6A deficiency (**Fig. 6J**). There are currently no treatment options for KAT6A Syndrome. Our study suggests that enhancing Wnt signaling in hippocampus is a potential therapeutic strategy for intellectual disability, a major clinical feature of this disease.

While histone acetyltransferases can potentially regulate the expression of many genes by altering the chromosome structure, it’s interesting that relatively few changes were detected in hippocampus after the loss of *Kat6a*. Only *Rspo2* stood out as a robust transcriptional target of KAT6A. This may reflect functional redundancy between various MYST family proteins, especially its paralog KAT6B. Indeed, only acetylation of H3K23 in *Rspo2* promoter region was reduced, while acetylation of other lysine residues was normal. Nevertheless, RSPO2 appears to be the key mediator downstream of KAT6A in hippocampus. First, RSPO2 deficiency phenocopies the loss of *Kat6a*, impairing CA3 synaptic functions and causing memory defects. Second, overexpressing RSPO2 in CA3 pyramidal neurons largely rescues the deficits in both Wnt signaling and memory-related behaviors in *Kat6a* mutant mice. These results support a crucial role of KAT6A-RSPO2 axis in the regulation of synaptic plasticity and hippocampus-dependent memory, and underscore the importance of Wnt signaling in the cognitive functions. Notably, *Rspo2* is also highly enriched in basolateral amygdala (BLA) excitatory neurons and functions as a neuronal marker for negative behaviors (*28*). Cre recombinase in *NEX*-cre line was not detected in BLA neurons, suggesting that memory deficits caused by the deletion of KAT6A or RSPO2 in excitatory neurons are unlikely due to the downregulation of RSPO2 expression in BLA. In addition, osteoblast-specific *Rspo2* knockout mice displayed decreased body size and bone mass (*29*), indicating the growth retardation associated with *Kat6a* haploinsufficiency may be attributable to the reduction of *Rspo2* in the osteoblasts. Taken together, given the importance of Wnt signaling, KAT6A/RSPO2/Wnt pathway may also play a role in other brain regions or peripheral organs, which are involved in the pathogenesis of KAT6A Syndrome.

Hippocampal CA1 region is important for learning and memory, and its neuronal defects are widely implicated in ID-related neurodevelopmental diseases (*30–32*). Surprisingly, loss of KAT6A did not affect CA1 synaptic function. Instead, it impaired CA3 pyramidal neuron synaptic transmission and plasticity, especially those of synapses receiving the prominent recurrent collateral inputs. Consistent with the functional deficits, KAT6A deficiency also decreased dendritic spine density at stratum radiatum layer of CA3 pyramidal neurons, the main region for CA3 recurrent collateral connections. All these defects are dependent on KAT6A-mediated downregulation of CA3-enriched *Rspo2*, suggesting a unique requirement of strong Wnt signaling for the development and function of CA3 pyramidal neurons. Although Wnt signaling has been involved in regulating synaptic function in other brain regions (*33, 34*), its molecular mechanism and downstream target genes are still poorly understood. The important role of Wnt signaling in CA3 as we identify here may provide a good model to fill this gap in the future. Altogether, our work links CA3 dysfunctions and Wnt signaling defects to cognitive deficits associated with KAT6A Syndrome, which may also have implications to other ID-related diseases.

## MATERIALS AND METHODS

### Animals

All procedures related to animal care and treatment were approved by the Johns Hopkins University Animal Care and Use Committee and met the guidelines of the National Institute of Health Guide for the Care and Use of Laboratory Animals. All animals were group housed in a standard 12-hour light/dark cycle with ad libitum access to food and water. Male mice aged 2-3 months were used for all experiments unless otherwise noted. Female mice were also used for some of the tests. The following mouse lines (8-12 weeks old) were used for the experiments: C57BL/6J (Jackson Laboratory, 000664), *Camk2a*-cre (Jackson Laboratory, 005359), *Gfap*-cre (Jackson Laboratory, 024098), *Gad2*-Cre (Jackson Laboratory, 010802), and Ai14 (B6.Cg-Gt(*ROSA*)26*Sor^tm14(CAG-tdTomato)Hze^*/J, Jackson Laboratory, 007914). *NEX*-cre was a gift from Klaus-Armin Nave (*35*). *Rspo2* floxed mice were kindly provided from Dr. Kurt Hankenson’s laboratory at University of Michigan (*29*). The sequences of the primers used for PCR genotyping are provided in **Table S2**.

### Cell culture and siRNA transfection

Human THP-1 cells used were purchased from ATCC and maintained in RPMI-1640 medium supplemented with 10% fetal bovine serum (FBS) and 1% Non-Essential Amino Acid (NEAA) at 37 °C in humidified 95% CO_2_ incubators. Cells were tested negative for mycoplasma contamination before use. siRNAs were purchased from Dharmacon and transfected into THP-1 cells with the final concentration of 30 nM using Lipofectamine RNAiMAX (Invitrogen) according to the manufacturer’s instructions. The sequences of siRNA used in this study are as follows: siControl sense: UGGUUUACAUGUCGACUAA, KAT6A siRNA#1 sense: GGAGUUGAGUGUUAAAGAU, siRNA#2 sense: GCGCUAUACUAAUCCAAUA.

### Generation of *Kat6a* floxed mice

*Kat6a* floxed mice were generated at Transgenic Core of Johns Hopkins University using the *Easi*-CRISPR (Efficient additions with ssDNA inserts-CRISPR) method, as previously described (*14*). Two single-guide RNAs (sgRNAs) were designed by http://crispor.tefor.net/. The sequences were as follows: sgRNA #1 (reverse strand): ATCTGGGATCTTATCATATCTGG; sgRNA #2 (forward strand): TTGAGCTGTTTATACTACAGAGG. Two crRNAs containing each sgRNA and ssDNA donor containing the homology arms and the floxed exon sequences were custom synthesized from IDT company. The annealed crRNA and tracrRNA (IDT) were diluted in microinjection buffer (0.25 mM EDTA, 10 mM Tris-HCl, pH 7.4) and mixed with Cas9 protein (30 ng/µl, IDT) to obtain ctRNP complexes. One-cell embryos of C57BL/6J mice were microinjected with a mixture of floxing ssDNA donors and two ctRNP complexes, and were transferred into the oviducts of pseudopregnant ICR females (Charles River). Successful insertions of two LoxP sites were detected by PCR genotyping of mouse tails and confirmed by Sanger sequencing (**Fig. S1A**). During the generation of mice with floxed exons, *Kat6a* knockout mice were also obtained due to the failed homology-directed repair at Cas9 cleavage sites, which were verified by PCR genotyping and Sanger sequencing (**Fig. S1B-C**). The sequences of the primers used are provided in **Table S2**.

### Generation of *Kat6a* 3**×**HA knockin mice

*Kat6a* 3×HA knockin mice were generated at Transgenic Core of Johns Hopkins University using the *Easi*-CRISPR method. One single-guide RNAs (sgRNAs) were designed by http://crispor.tefor.net/. The sequence was as follows: sgRNA (forward strand): CTTACATGAGGAGATGAGCGAGG. One crRNAs containing sgRNA and ssDNA donor containing the homology arms and the 3×HA sequences were custom synthesized from IDT company. The annealed crRNA and tracrRNA (IDT) were diluted in microinjection buffer (0.25 mM EDTA, 10 mM Tris-HCl, pH 7.4) and mixed with Cas9 protein (30 ng/µl, IDT) to obtain ctRNP complexes. One-cell embryos of B6SJLF2 mice (Jackson Lab) were microinjected with a mixture of ssDNA donors and the ctRNP complexes, and were transferred into the oviducts of pseudopregnant ICR females (Charles River). Successful insertions of 3×HA were detected by PCR genotyping of mouse tails and confirmed by Sanger sequencing. The sequences of the primers used are provided in **Table S2**.

### Real-time qPCR

Total RNA was isolated from samples with Trizol reagents (Invitrogen) and any potential DNA contamination was removed by RNase-free DNase treatment (Promega, M6101). mRNA was reverse-transcribed into cDNA using the high-capacity cDNA reverse transcription kits (Applied Biosystems, #4368814). Relative quantitation was determined using the QuantStudio 6 Flex detection system (Applied Biosystems) that measures real-time SYBR green fluorescence and then calculated by means of the comparative Ct method (2^−ΔΔCt^) with the expression of *Gapdh* as an internal control. The sequences of the primers used are provided in **Table S3**.

### Western blot

Proteins was isolated from cultured cells or brain tissues with RIPA buffer (Sigma, R0278) including protease inhibitors cocktails. Samples were resolved on SDS/PAGE and transferred to nitrocellulose membranes (Bio-Rad, #1620115), which were incubated with appropriate antibodies for overnight at 4 °C. Primary antibody concentrations: anti-HA (rabbit, 1:1000, Cell Signaling, #3724), anti-β-actin (mouse, 1:5000, Proteintech, 66009-1-lg), anti-FLAG (mouse, 1:1000, Sigma, F1804), anti-non-phospho (active) β-catenin (rabbit, 1:1000, Cell Signaling, #8814), anti-KAT6A (rabbit, 1:1000, Invitrogen, PA5_68046), anti-RSPO2 (rabbit, 1:1000, Biorbyt, orb185986), anti-JNK (rabbit, 1:1000, Cell Signaling, #9252), anti-phospho-JNK (rabbit, 1:1000, Cell Signaling, #9251). After wash, the membranes were incubated a HRP-conjugated secondary antibody (Cytiva, 1:5000). Immunoreactive bands were visualized using Western Chemiluminescent HRP Substrate (Millipore, #WBKLS0500) and analyzed with ImageJ (NIH).

### Immunohistochemistry

Anesthetized mice were perfused transcardially with PBS, followed by 4% cold paraformaldehyde (PFA) in PBS. Brains were removed and post-fixed in 4% PFA at 4 °C overnight. After dehydration by 30% sucrose, brains were embedded in OCT (Tissue-Tek) and cut into 30-µm-thick sections on cryostat microtome (Leica). Sections were permeabilized with 0.3% Triton X-100 and 5% BSA in PBS for 1 h at room temperature, washed with PBS three times, blocked in 10% BSA, and incubated with primary antibodies at 4 °C overnight. Primary antibody concentrations: anti-FLAG (mouse, 1:100, Sigma, F1804), anti-DCX (rabbit, 1:100, Cell Signaling, #4604), anti-BrdU (mouse, 1:100, Bio-Rad, MCA2483GA), anti-NeuN (rabbit, 1:200, Cell Signaling, #12943). After washing 3 times with PBS, samples were incubated with Alexa Fluor-conjugated secondary antibodies (1:500, Invitrogen) for 1-2 h at RT. Fluorescent images were taken using a confocal microscope (Zeiss LSM 900) and analyzed with ImageJ software (NIH).

### Chromatin immunoprecipitation qPCR

Chromatin immunoprecipitation (ChIP) experiments were performed according to the procedure described previously (*36*). Hippocampus tissues from 3 mice were fixed with 1% formaldehyde for 15 min at room temperature. The fixed cells were lysed in lysis buffer (5 mM EDTA, 50 mM Tris-HCl (pH 8.1) and 1% SDS, plus protease inhibitor cocktail). The lysates were then sonicated with 3 × 10 cycles (30 s on and off) (Bioruptor, Diagenode) to generate chromatin fragments of ∼500 bp in length. Cell debris was removed by centrifugation and supernatant were collected. A dilution buffer (2 mM EDTA, 1% Triton X-100, 150 mM NaCl, 20 mM Tris-HCl (pH 8.1), plus protease inhibitor cocktail) was subsequently applied (1:10 ratio) and the resultant chromatin solution (aliquot 50 µl as the input) was then incubated with control or specific antibodies (3-5 µg) for 12 h at 4 °C with constant rotation. Protein A/G Sepharose beads (50 µl of 50% (vol/vol)) were added for incubation of another 2 h. Beads were collected by centrifugation at 500 g for 5 min at 4 °C. Beads were sequentially washed with the following buffers for 5 min at 4 °C: TSE I (0.1% SDS, 1% Triton X-100, 2 mM EDTA, 20 mM Tris-HCl (pH 8.1), 150 mM NaCl); TSE II (0.1% SDS, 2 mM EDTA, 1% Triton X-100, 20 mM Tris-HCl (pH 8.1), 500 mM NaCl); buffer III (0.25 M LiCl, 1% Nonidet P-40, 1 mM EDTA, 1% sodium deoxycholate and 10 mM Tris-HCl (pH 8.1)); Tris-EDTA buffer. The input and the precipitated DNA-protein complex were decrosslinked at 65 °C for 12 h in elution buffer (5 mM EDTA, 1% SDS, 50 mM NaCl, 20 mM Tris-HCl (pH 8.1), 0.1 mg/ml proteinase K), and DNA was purified using PCR purification kit (Qiagen). Quantification of the precipitated DNA fragments were performed with real-time PCR using primers listed in **Table S4**.

### RNA sequencing

Four biological replicates were sequenced per group. For each sample, RNA was extracted from hippocampus tissues of littermate controls and *Kat6a*^+/-^ mice, followed by purification with the RNeasy Micro kit (QIAGEN, #74004). High-throughput RNA sequencing (RNA-seq) was performed by Illumina NovaSeq 6000 at Novogene (CA, USA). The raw sequencing data were aligned to the mouse preference genome (GRCm38, mm10) using STAR software (v2.5). Reads on each GENCODE annotated gene were counted using HTSeq, and then differential gene expression analysis was performed using DESeq2 R package.

### Nuclei isolation

Mouse hippocampal tissues were rapidly dissected in ice-cold homogenization buffer (0.25 M sucrose, 25 mM KCl, 5 mM MgCl_2_, 20 mM Tricine-KOH, pH=7.8). The tissue was homogenized using a loose-fitting pestle in 1.5 ml of homogenization buffer supplemented with 1 mM DTT, 0.15 mM spermine, 0.5 mM spermidine, 1×EDTA-free protease inhibitor (Roche), and 60 U/mL RNasin Plus RNase Inhibitor (Promega, N2611). A 6% IGEPAL CA-630 solution was added to bring the homogenate to 0.3% IGEPAL CA-630, and the homogenate was further homogenized with five strokes of a tight-fitting pestle. The sample was filtered through a 50 µm filter (Sysmex, 04-004-2327), underlayed with solutions of 30% and 40% iodixanol (Sigma, D1556) in homogenization buffer, and centrifuged at 10,000 g for 20 min in a swinging bucket centrifuge at 4 °C. Nuclei were collected at the 30-40% interface, diluted with two volumes of homogenization buffer and concentrated by centrifugation for 10 min at 500 g at 4 °C. These nuclei can be used for sorting or snRNA sequencing.

### snRNA-seq

Two biological replicates were sequenced per genotype. snRNA-seq libraries were constructed using the 10X Genomics Chromium single cell kit. Libraries were sequenced on an Illumina NovaSeq 6000 at Single Cell & Transcriptomics Core of Johns Hopkins University. Reads were aligned to the mm10 reference genome using the kb-python program (version 0.25.0) (*37*). The data was then imported into R and low quality nuclei and “empty droplets” were filtered out using the barcodeRanks from the DropletUtils package. The data was analyzed using the Seurat R package (version 4.0.1) as described previously (*38*). Briefly, data normalization was performed using SCTransform function. The uwot package (https://github.com/jlmelville/uwot) was used for UMAP dimensional reduction. Seurat FindMarkers function was used for the analysis of differential gene expression and plotted using ggplot2 and the tidyverse collection of R packages in RStudio.

### RNAscope *in situ* hybridization

Fixed brains were embedded in OCT (Tissue-Tek) and sectioned at a thickness of 14 µm. RNAscope Multiplex Fluorescent Reagent Kit v2 (ACD, Advanced Cell Diagnostics) was used following the manufacturer’s manual for the fixed frozen tissues. Probe targeting mouse *Rspo2* (#402001) was purchased from ACD. TSA Plus fluorescein (#NEL741E001KT), or Cyanine 3 (#NEL744001KT) was used for developing the fluorescence signal. Images were collected with a Zeiss LSM 900 confocal microscope and analyzed using ImageJ software.

### Stereotaxic virus injection

Mice (3-6 weeks old) were anesthetized with the inhalation anesthetic isoflurane and positioned in the stereotaxic apparatus (RWD Life Sciences). After craniotomy, the mice were bilaterally injected with 0.5 µl AAV viruses into the CA3 (coordinates, bregma: anterior/posterior: −1.8 mm, medial/lateral: ±2.2 mm, dorsal/ventral: −2.3 mm). After the injection, the needle was maintained in the place for an additional 10 min to facilitate the diffusion of the viruses and then slowly withdrawn. The viruses used in this study were as follows: AAV5-CamkII-GFP-Cre purchased from UNC Vector Core, AAV5-CamkII-GFP purchased from Addgene, and AAV5-CamkII-RSPO2-FLAG packaged from WZ Biosciences Inc (Maryland, USA). All AAV viruses were diluted to titers of 3-5 ×10^12^ particles per ml. Mice receiving virus injections were returned to their home cages to recover for 4-5 weeks before they were used for experiments.

### Acute brain slice electrophysiology

For CA1 recordings, hippocampal transverse slices (300 µm for whole-cell recording while 400 µm for fEPSP recording) were cut from 6-8 weeks old mice using a Leica vibratome in chilled high-sucrose cutting solution containing (in mM): 2.5 KCl, 1 NaH_2_PO_4_, 26 NaHCO_3_, 7 glucose, 210 sucrose, 0.5 CaCl_2_ and 7 MgSO_4_. Slices were then incubated for 30 min at 34 °C in artificial cerebrospinal fluid (aCSF) bubbled with 95% O_2_ / 5% CO_2_ containing (in mM): 119 NaCl, 2.5 KCl, 26.2 NaHCO_3_, 1 NaH_2_PO_4_, 11 glucose, 2.5 CaCl_2_ and 1.3 MgSO_4_. Slices were recovered at room temperature for at least 1 hour. The slices were transferred to a submersion chamber on an upright Olympus microscope, and perfused in aCSF containing 0.1 mM picrotoxin and 0.01 mM bicuculline, saturated with 95% O_2_ / 5% CO_2_. The perfusion flow rate was maintained constant at 2 ml/min. Recordings were made with MultiClamp 700B amplifier and 1550B digitizer (Molecular Devices). Data acquisition were performed with pClamp 10.7 software (Molecular Devices), filtered at 2 kHz and digitized at 10 kHz. Whole-cell recordings were made by 3-5 MΩ borosilicate glass pipettes filled with intracellular solution consisted of (in mM): 135 CsMeSO_4_, 8 NaCl, 10 HEPES, 0.3 EGTA, 5 QX-314, 4 Mg-ATP, 0.3 Na-GTP and 0.1 spermine. mEPSCs were recorded at a holding potential of −70 mV in the presence of 1 µM TTX. The paired-pulse ratio was measured by applying two pulses at an interval of 25 ms, 50 ms, 100 ms, and 200 ms, respectively, and then calculating the ratio of the peak amplitudes. The input resistance and pipette series resistance were monitored throughout the recording. The field EPSPs (fEPSPs) were recorded via a glass micropipette filled with ACSF (1-3 MΩ) placed in the CA1 stratum radiatum. To induce LTP, the stimulation intensity was adjusted to evoke ∼30-50% of the maximum response. TBS-LTP was induced by four trains of theta burst stimulations (five pulses at 100 Hz every 200 ms) with an inter-train interval of 10 s. fEPSP slopes were measured by linear interpolation from 10-90% of maximum negative deflection.

For CA3 recordings, 6-8 weeks old mice were anesthetized with the inhalation anesthetic isoflurane, and then perfused intracardially with ice-cold oxygenated cutting solution containing (in mM): 110 choline chloride, 2.5 KCl, 1.25 NaH_2_PO_4_, 25 NaHCO_3_, 0.5 CaCl_2_, 7 MgCl_2_, 10 glucose, 3 Na-pyruvate, saturated with 95% O_2_ / 5% CO_2_. The brain was removed rapidly and immersed in ice-cold oxygenated cutting solution. Transverse hippocampal slices (300 µm for whole-cell recording while 400 µm for fEPSP recording) were cut in the cutting solution using a vibratome (VT-1200S, Leica) and transferred to artificial cerebrospinal fluid (aCSF) containing (in mM): 125 NaCl, 2.5 KCl, 1.25 NaH_2_PO_4_, 25 NaHCO_3_, 2 CaCl_2_, 2 MgCl_2_, 10 glucose, 3 Na-pyruvate, saturated with 95% O_2_ / 5% CO_2_. The slices were recovered for 30 min at 35 °C and then maintained at room temperature for ∼1 hour. Slices were subsequently transferred to a submerged recording chamber containing aCSF (without Na-pyruvate) perfusion maintained at 34 °C. Recordings were made with MultiClamp 700B amplifier (Molecular Devices). Whole-cell recordings were made by 3-5 MΩ borosilicate glass pipettes filled with intracellular solution consisted of (in mM): 120 K-gluconate, 15 KCl, 10 HEPES, 2 MgCl_2_, 0.2 EGTA, 4 Na_2_ATP, 0.3 Na_3_-GTP, 14 Tris-phosphocreatine (pH 7.3). Hippocampal CA3 pyramidal neurons were held at −70 mV and mEPSCs were recorded in the presence of 0.1 mM picrotoxin and 0.5 µM TTX. In the mossy fiber blockade experiments, mEPSCs were recorded with 1 µM DCG-IV in the aCSF. Series resistance was monitored throughout each experiment and controlled below 20 MΩ. Data was discarded if the series resistance varied more than 20%. For A/C-LTP recordings, a concentric bipolar electrode was placed in CA3 stratum radiatum to stimulate A/C synapses and field EPSPs (fEPSPs) were recorded with a glass pipette (2-3 MΩ) filled with aCSF. 0.1 mM picrotoxin was added to block inhibitory transmission and 1 µM DCG-IV was added to block mossy fiber inputs. The stimulus intensity was adjusted to evoke 40%-50% of the maximal response. LTP was induced by four trains of 100 Hz (1 s) tetanus with an inter-train interval of 10 s.

### Sholl analysis and spine analysis

Recorded pyramidal neurons of CA3 were filled with 0.2% (w/v) biocytin. Then, the slices containing the fixed neurons cross-linked with 4% paraformaldehyde and stained with streptavidin Alexa Fluor 555 conjugate (Invitrogen, 1:1000). Fluorescent images were obtained using a confocal microscope (ZEISS LSM 900) and imported into ImageJ (NIH) for further analysis. For Sholl analysis, concentric circles at 20 µm intervals were drawn around the soma. The number of dendrites crossing each circle was calculated and the data were presented as mean ± s.e.m. For spine analysis, dendritic segments at stratum radiatum layer of CA3 were randomly chosen and images were taken with a resolution of 1024 × 1024. An average of three dendrites per neuron on 4 neurons per mouse (n = 12 dendrites per mice, n = 4 mice per group) totaling ∼2,000 dendritic spines per experimental group were analyzed. Spines with a neck can be classified as either thin or mushroom, and those without a significant neck are classified as stubby (*39*). Spines with a neck were labeled as thin or mushroom based on head diameter. These parameters have been verified by comparison with trained human operators. The number of dendrites spines is presented as mean ± s.e.m.

### Behavioral tests

Mice were handled by investigators for four days before any behavioral test. Mice were given a 2-week interval to recover from the first behavioral test before the next behavioral test. Mice were not subjected to more than three behavioral tests. All behavioral tests were performed at Behavioral Core of Johns Hopkins University.

In the open field test, mice were placed in a chamber (18” × 18”) with infrared beams (Photobeam activity system, San Diego Instruments) and monitored for movement by using horizontal photobeams. Beam breaks were converted to directionally specific movements and summated at 5-min intervals over 15 min. Ambulatory activity (beam breaks) was measured as total horizontal photobeam breaks.

In the elevated plus maze test, we used a plus-shaped platform at 40 cm above the floor with with two open arms (30 cm × 5 cm) and two closed arms (30 cm × 5 cm × 15 cm) on opposing sides of a central square platform (5 cm × 5 cm). The overall illuminations of the 4 arms were kept equal, at 100-200 lux. Each animal was gently placed in the center platform facing an open arm and was videotaped for 5 min. The total time and entries in the open arms and the distance the animals traveled were measured using the ANY-maze software. The number of visits to the open arms and the time spent on the open arms were used as measures for anxiety.

For the social interaction test, we used a three-chamber apparatus (60 × 40 × 22 cm) that consisted of three chambers (left, center, and right) of equal size with 10 × 5 cm openings between the chambers. The test consisted of two phases with different stimulus in each of two side chambers. The stimulus was placed inside a capsule (an inverted mesh pencil cup). During habituation, mice were allowed to explore the apparatus with two empty capsules for 10 minutes. During testing, the first phase contained two identical non-social stimuli (paper ball) and the second phase contained a non-social stimulus (a Lego cube) and a social stimulus (unfamiliar mouse matched in strain, sex and age). The test animal was placed in the center of the chamber and allowed to explore the apparatus in the presence of these stimuli for 10 minutes. Animal behavior during the test session was tracked by a top camera, analyzed by ANY-maze software and verified manually. Interaction time was counted based on close “investigating” behaviors of the test animal to each stimulus. Social preference index = (social time - non-social time) / (social time + non-social time). The partner mice were used up to three times, with one test per day.

In the novel object recognition test, we used a small chamber (30 cm × 30 cm) to facilitate object exploration and reduce the time needed to habituate mice to the arena. The arena was surrounded on three sides by a white screen to limit spatial information and prevent spatial biases. During the training session, two identical objects (Lego) were placed at left and right sides of the arena. During the training, mice were placed in the center of the arena and allowed to explore freely for 10 min. Mice were returned to their home cages. Then 2 h later, one of the objects was replaced by a novel object with different color and shape, and mice were allowed to freely explore the whole arena for 5 min. Animal behavior during the test session was tracked by a top camera, analyzed by ANY-maze software and verified manually. The discrimination index was calculated as (exploration time on the novel object - exploration time on the familiar object) / (exploration time on the novel object + exploration time on the familiar object). Sniffing, touching (> 1 s), and staring the objects were judged as exploration behaviors.

In the Morris water maze test, we used a maze consisted of a round pool (diameter: 120 cm) filled with water that was at 24 °C and made opaque with non-toxic white tempera paint. A circular plastic platform (diameter: 10 cm) was placed at the center of the target quadrant and submerged 1 cm below the surface of the water. Four local cues were provided to allow spatial map generation. The Morris water maze test was performed as previously described (*40*). Briefly, we trained the mice for 4 trials per day with different start points for 5 consecutive days. Mice were gently placed into the water facing the wall of the pool and allowed to freely explore the whole maze for 1 min. Mice were then guided to the rescue platform if they did not find it. Mice were allowed to take a rest on the platform for 10 s, and then re-trained from a different start position with the same procedure. The latency for each animal to find the platform (at least 3 s stay) was recorded. On Day 6, the platform was removed, and animals searched freely for 1 min starting from the opposite quadrant. The entries into the platform area, total time spent in the target quadrant, and the total distance travels were recorded using the ANY-maze software. The animals with an average speed below 10 cm/sec were excluded for their weak motor ability.

In the fear conditioning test, mice were first habituated to the conditioning chamber (Med Accociates Inc) for 10 min the day before training. On the day of training, after 3 min exploration in the conditioning chamber, each mouse received one pairing of a tone (2,800 Hz, 80 dB, 30 s) with a short co-terminating foot shock (0.5 mA, 2 s), after which they remained in the chamber for additional 30 s and were then returned to home cages. 24 h after the conditioning, mice were tested for freezing (behavioral immobility) in response to the training context (training chamber) and to the tone (in the training chamber with a new environment and odour). The percentage of freezing time was calculated as an index of fear learning and memory. For contextual fear memory tests, mice were returned to the conditioning chamber for 3 min and freezing behavior was counted using Video Freeze software (Med Accociates Inc). For auditory fear memory tests, mice were placed in a changed chamber and freezing responses were recorded during the last 3 min when the tone was delivered.

### Statistical analysis

Statistical analyses were performed using GraphPad Prism 9.4 (GraphPad Software, La Jolla, CA). Before statistical analysis, variation within each group of data and the assumptions of the tests were checked. For in vivo experiments, the animals were distributed into various treatment groups randomly. For in vitro experiments, the cells were evenly suspended and then randomly distributed in each well tested. Comparisons between two groups were made using unpaired Student’s two-tailed t test or Mann-Whitney test. Comparisons among three or more groups were made using one- or two-way ANOVA followed by Bonferroni’s post hoc test. All experiments and analysis of data were performed in a blinded manner by investigators who were unaware of the genotype or manipulation. The significance level was set at p < 0.05. Test statistics, n numbers, and p values are indicated in the figure legends. All data are presented as mean ± SEM.

## Acknowledgments

We thank Dr. Kurt Hankenson for kindly providing the *Rspo2*-floxed mice, Dr. Amir Rattner and Dr. Chaohua Jiang for their technical assistance to this project, and Dr. Jeremy Nathans and members of the Qiu lab for valuable discussions.

## Author contributions

Y.L. initiated the project and performed the majority of the experiments. M.F. and J.Y. performed electrophysiological recordings. L.M., Y.Y. and S.S. analyzed the sequencing data. K.H.C. performed the genotyping of *Rspo2* mice. Y.L., M.F., J.Y., L.M., and Z.Q. analyzed and interpreted the results. Y.L. and Z.Q. designed the study and wrote the paper with input from all authors.

## Data and materials availability

RNA-seq data and snRNA-seq data have been deposited in Gene Expression Omnibus (GEO) with accession number of GSE261058 and GSE261148, respectively. All other data needed to evaluate the conclusions in the paper are present in the paper and/or the Supplementary Materials.

## REFERENCES

1. E. R. Kandel, Y. Dudai, M. R. Mayford, The molecular and systems biology of memory. Cell 157, 163–186 (2014).

2. E. L. Yap, M. E. Greenberg, Activity-Regulated Transcription: Bridging the Gap between Neural Activity and Behavior. Neuron 100, 330–348 (2018).

3. K. K. Lee, J. L. Workman, Histone acetyltransferase complexes: one size doesn’t fit all. Nat Rev Mol Cell Biol 8, 284–295 (2007).

4. J. Humbert, S. Salian, P. Makrythanasis, G. Lemire, J. Rousseau, S. Ehresmann, T. Garcia, R. Alasiri, A. Bottani, S. Hanquinet, E. Beaver, J. Heeley, A. C. M. Smith, S. I. Berger, S. E. Antonarakis, X. J. Yang, J. Cote, P. M. Campeau, De Novo KAT5 Variants Cause a Syndrome with Recognizable Facial Dysmorphisms, Cerebellar Atrophy, Sleep Disturbance, and Epilepsy. Am J Hum Genet 107, 564–574 (2020).

5. E. Tham, A. Lindstrand, A. Santani, H. Malmgren, A. Nesbitt, H. A. Dubbs, E. H. Zackai, M. J. Parker, F. Millan, K. Rosenbaum, G. N. Wilson, A. Nordgren, Dominant mutations in KAT6A cause intellectual disability with recognizable syndromic features. Am J Hum Genet 96, 507–513 (2015).

6. K. Yan, J. Rousseau, R. O. Littlejohn, C. Kiss, A. Lehman, J. A. Rosenfeld, C. T. R. Stumpel, A. P. A. Stegmann, L. Robak, F. Scaglia, T. T. M. Nguyen, H. Fu, N. F. Ajeawung, M. V. Camurri, L. Li, A. Gardham, B. Panis, M. Almannai, M. J. G. Sacoto, B. Baskin, C. Ruivenkamp, F. Xia, W. Bi, D. D. D. Study, C. Study, M. T. Cho, T. P. Potjer, G. W. E. Santen, M. J. Parker, N. Canham, M. McKinnon, L. Potocki, J. J. MacKenzie, E. R. Roeder, P. M. Campeau, X. J. Yang, Mutations in the Chromatin Regulator Gene BRPF1 Cause Syndromic Intellectual Disability and Deficient Histone Acetylation. Am J Hum Genet 100, 91–104 (2017).

7. L. Li, M. Ghorbani, M. Weisz-Hubshman, J. Rousseau, I. Thiffault, R. E. Schnur, C. Breen, R. Oegema, M. M. Weiss, Q. Waisfisz, S. Welner, H. Kingston, J. A. Hills, E. M. Boon, L. Basel-Salmon, O. Konen, H. Goldberg-Stern, L. Bazak, S. Tzur, J. Jin, X. Bi, M. Bruccoleri, K. McWalter, M. T. Cho, M. Scarano, G. B. Schaefer, S. S. Brooks, S. S. Hughes, K. L. I. van Gassen, J. M. van Hagen, T. K. Pandita, P. B. Agrawal, P. M. Campeau, X. J. Yang, Lysine acetyltransferase 8 is involved in cerebral development and syndromic intellectual disability. J Clin Invest 130, 1431–1445 (2020).

8. J. Clayton-Smith, J. O’Sullivan, S. Daly, S. Bhaskar, R. Day, B. Anderson, A. K. Voss, T. Thomas, L. G. Biesecker, P. Smith, A. Fryer, K. E. Chandler, B. Kerr, M. Tassabehji, S. A. Lynch, M. Krajewska-Walasek, S. McKee, J. Smith, E. Sweeney, S. Mansour, S. Mohammed, D. Donnai, G. Black, Whole-exome-sequencing identifies mutations in histone acetyltransferase gene KAT6B in individuals with the Say-Barber-Biesecker variant of Ohdo syndrome. Am J Hum Genet 89, 675–681 (2011).

9. J. Borrow, V. P. Stanton, Jr., J. M. Andresen, R. Becher, F. G. Behm, R. S. Chaganti, C. I. Civin, C. Disteche, I. Dube, A. M. Frischauf, D. Horsman, F. Mitelman, S. Volinia, A. E. Watmore, D. E. Housman, The translocation t(8;16)(p11;p13) of acute myeloid leukaemia fuses a putative acetyltransferase to the CREB-binding protein. Nat Genet 14, 33–41 (1996).

10. J. B. Baell, D. J. Leaver, S. J. Hermans, G. L. Kelly, M. S. Brennan, N. L. Downer, N. Nguyen, J. Wichmann, H. M. McRae, Y. Yang, B. Cleary, H. R. Lagiakos, S. Mieruszynski, G. Pacini, H. K. Vanyai, M. I. Bergamasco, R. E. May, B. K. Davey, K. J. Morgan, A. J. Sealey, B. Wang, N. Zamudio, S. Wilcox, A. L. Garnham, B. N. Sheikh, B. J. Aubrey, K. Doggett, M. C. Chung, M. de Silva, J. Bentley, P. Pilling, M. Hattarki, O. Dolezal, M. L. Dennis, H. Falk, B. Ren, S. A. Charman, K. L. White, J. Rautela, A. Newbold, E. D. Hawkins, R. W. Johnstone, N. D. Huntington, T. S. Peat, J. K. Heath, A. Strasser, M. W. Parker, G. K. Smyth, I. P. Street, B. J. Monahan, A. K. Voss, T. Thomas, Inhibitors of histone acetyltransferases KAT6A/B induce senescence and arrest tumour growth. Nature 560, 253–257 (2018).

11. B. N. Sheikh, N. L. Downer, B. Phipson, H. K. Vanyai, A. J. Kueh, D. J. McCarthy, G. K. Smyth, T. Thomas, A. K. Voss, MOZ and BMI1 play opposing roles during Hox gene activation in ES cells and in body segment identity specification in vivo. Proc Natl Acad Sci U S A 112, 5437–5442 (2015).

12. T. Katsumoto, Y. Aikawa, A. Iwama, S. Ueda, H. Ichikawa, T. Ochiya, I. Kitabayashi, MOZ is essential for maintenance of hematopoietic stem cells. Genes Dev 20, 1321–1330 (2006).

13. J. Kennedy, D. Goudie, E. Blair, K. Chandler, S. Joss, V. McKay, A. Green, R. Armstrong, M. Lees, B. Kamien, B. Hopper, T. Y. Tan, P. Yap, Z. Stark, N. Okamoto, N. Miyake, N. Matsumoto, E. Macnamara, J. L. Murphy, E. McCormick, H. Hakonarson, M. J. Falk, D. Li, P. Blackburn, E. Klee, D. Babovic-Vuksanovic, S. Schelley, L. Hudgins, S. Kant, B. Isidor, B. Cogne, K. Bradbury, M. Williams, C. Patel, H. Heussler, C. Duff-Farrier, P. Lakeman, I. Scurr, U. Kini, M. Elting, M. Reijnders, J. Schuurs-Hoeijmakers, M. Wafik, A. Blomhoff, C. A. L. Ruivenkamp, E. Nibbeling, A. J. M. Dingemans, E. D. Douine, S. F. Nelson, D. D. D. Study, M. Hempel, T. Bierhals, D. Lessel, J. Johannsen, V. A. Arboleda, R. Newbury-Ecob, KAT6A Syndrome: genotype-phenotype correlation in 76 patients with pathogenic KAT6A variants. Genet Med 21, 850–860 (2019).

14. H. Miura, R. M. Quadros, C. B. Gurumurthy, M. Ohtsuka, Easi-CRISPR for creating knock-in and conditional knockout mouse models using long ssDNA donors. Nat Protoc 13, 195–215 (2018).

15. I. Izquierdo, C. R. Furini, J. C. Myskiw, Fear Memory. Physiol Rev 96, 695–750 (2016).

16. H. Eichenbaum, Hippocampus: cognitive processes and neural representations that underlie declarative memory. Neuron 44, 109–120 (2004).

17. T. Nakashiba, J. Z. Young, T. J. McHugh, D. L. Buhl, S. Tonegawa, Transgenic inhibition of synaptic transmission reveals role of CA3 output in hippocampal learning. Science 319, 1260–1264 (2008).

18. K. Nakazawa, M. C. Quirk, R. A. Chitwood, M. Watanabe, M. F. Yeckel, L. D. Sun, A. Kato, C. A. Carr, D. Johnston, M. A. Wilson, S. Tonegawa, Requirement for hippocampal CA3 NMDA receptors in associative memory recall. Science 297, 211–218 (2002).

19. T. A. Macek, D. G. Winder, R. W. t. Gereau, C. O. Ladd, P. J. Conn, Differential involvement of group II and group III mGluRs as autoreceptors at lateral and medial perforant path synapses. J Neurophysiol 76, 3798–3806 (1996).

20. H. Kamiya, S. Ozawa, Dual mechanism for presynaptic modulation by axonal metabotropic glutamate receptor at the mouse mossy fibre-CA3 synapse. J Physiol 518 (Pt 2), 497–506 (1999).

21. O. Kazanskaya, A. Glinka, I. del Barco Barrantes, P. Stannek, C. Niehrs, W. Wu, R-Spondin2 is a secreted activator of Wnt/beta-catenin signaling and is required for Xenopus myogenesis. Dev Cell 7, 525–534 (2004).

22. S. Seshagiri, E. W. Stawiski, S. Durinck, Z. Modrusan, E. E. Storm, C. B. Conboy, S. Chaudhuri, Y. Guan, V. Janakiraman, B. S. Jaiswal, J. Guillory, C. Ha, G. J. Dijkgraaf, J. Stinson, F. Gnad, M. A. Huntley, J. D. Degenhardt, P. M. Haverty, R. Bourgon, W. Wang, H. Koeppen, R. Gentleman, T. K. Starr, Z. Zhang, D. A. Largaespada, T. D. Wu, F. J. de Sauvage, Recurrent R-spondin fusions in colon cancer. Nature 488, 660–664 (2012).

23. E. Szenker-Ravi, U. Altunoglu, M. Leushacke, C. Bosso-Lefevre, M. Khatoo, H. Thi Tran, T. Naert, R. Noelanders, A. Hajamohideen, C. Beneteau, S. B. de Sousa, B. Karaman, X. Latypova, S. Basaran, E. B. Yucel, T. T. Tan, L. Vlaminck, S. S. Nayak, A. Shukla, K. M. Girisha, C. Le Caignec, N. Soshnikova, Z. O. Uyguner, K. Vleminckx, N. Barker, H. Kayserili, B. Reversade, RSPO2 inhibition of RNF43 and ZNRF3 governs limb development independently of LGR4/5/6. Nature 557, 564–569 (2018).

24. D. Lv, F. Jia, Y. Hou, Y. Sang, A. A. Alvarez, W. Zhang, W. Q. Gao, B. Hu, S. Y. Cheng, J. Ge, Y. Li, H. Feng, Histone Acetyltransferase KAT6A Upregulates PI3K/AKT Signaling through TRIM24 Binding. Cancer Res 77, 6190–6201 (2017).

25. B. J. Klein, S. M. Jang, C. Lachance, W. Mi, J. Lyu, S. Sakuraba, K. Krajewski, W. W. Wang, S. Sidoli, J. Liu, Y. Zhang, X. Wang, B. M. Warfield, A. J. Kueh, A. K. Voss, T. Thomas, B. A. Garcia, W. R. Liu, B. D. Strahl, H. Kono, W. Li, X. Shi, J. Cote, T. G. Kutateladze, Histone H3K23-specific acetylation by MORF is coupled to H3K14 acylation. Nat Commun 10, 4724 (2019).

26. H. X. Hao, Y. Xie, Y. Zhang, O. Charlat, E. Oster, M. Avello, H. Lei, C. Mickanin, D. Liu, H. Ruffner, X. Mao, Q. Ma, R. Zamponi, T. Bouwmeester, P. M. Finan, M. W. Kirschner, J. A. Porter, F. C. Serluca, F. Cong, ZNRF3 promotes Wnt receptor turnover in an R-spondin-sensitive manner. Nature 485, 195–200 (2012).

27. R. Sun, L. He, H. Lee, A. Glinka, C. Andresen, D. Hubschmann, I. Jeremias, K. Muller-Decker, C. Pabst, C. Niehrs, RSPO2 inhibits BMP signaling to promote self-renewal in acute myeloid leukemia. Cell Rep 36, 109559 (2021).

28. J. Kim, M. Pignatelli, S. Xu, S. Itohara, S. Tonegawa, Antagonistic negative and positive neurons of the basolateral amygdala. Nat Neurosci 19, 1636–1646 (2016).

29. M. N. Knight, K. Karuppaiah, M. Lowe, S. Mohanty, R. L. Zondervan, S. Bell, J. Ahn, K. D. Hankenson, R-spondin-2 is a Wnt agonist that regulates osteoblast activity and bone mass. Bone Res 6, 24 (2018).

30. G. L. Caldeira, A. S. Inacio, N. Beltrao, C. A. V. Barreto, M. V. Rodrigues, T. Rondao, R. Macedo, R. P. Gouveia, M. Edfawy, J. Guedes, B. Cruz, S. R. Louros, I. S. Moreira, J. Peca, A. L. Carvalho, Aberrant hippocampal transmission and behavior in mice with a stargazin mutation linked to intellectual disability. Mol Psychiatry 27, 2457–2469 (2022).

31. A. Pavlowsky, J. Chelly, P. Billuart, Emerging major synaptic signaling pathways involved in intellectual disability. Mol Psychiatry 17, 682–693 (2012).

32. S. Jawaid, G. J. Kidd, J. Wang, C. Swetlik, R. Dutta, B. D. Trapp, Alterations in CA1 hippocampal synapses in a mouse model of fragile X syndrome. Glia 66, 789–800 (2018).

33. A. Marzo, S. Galli, D. Lopes, F. McLeod, M. Podpolny, M. Segovia-Roldan, L. Ciani, S. Purro, F. Cacucci, A. Gibb, P. C. Salinas, Reversal of Synapse Degeneration by Restoring Wnt Signaling in the Adult Hippocampus. Curr Biol 26, 2551–2561 (2016).

34. R. F. Narvaes, C. R. G. Furini, Role of Wnt signaling in synaptic plasticity and memory. Neurobiol Learn Mem 187, 107558 (2022).

35. S. Goebbels, I. Bormuth, U. Bode, O. Hermanson, M. H. Schwab, K. A. Nave, Genetic targeting of principal neurons in neocortex and hippocampus of NEX-Cre mice. Genesis 44, 611–621 (2006).

36. Y. Liu, S. Lai, W. Ma, W. Ke, C. Zhang, S. Liu, Y. Zhang, F. Pei, S. Li, M. Yi, Y. Shu, Y. Shang, J. Liang, Z. Huang, CDYL suppresses epileptogenesis in mice through repression of axonal Nav1.6 sodium channel expression. Nat Commun 8, 355 (2017).

37. P. Melsted, A. S. Booeshaghi, L. Liu, F. Gao, L. Lu, K. H. J. Min, E. da Veiga Beltrame, K. E. Hjorleifsson, J. Gehring, L. Pachter, Modular, efficient and constant-memory single-cell RNA-seq preprocessing. Nat Biotechnol 39, 813–818 (2021).

38. J. Wang, A. Rattner, J. Nathans, A transcriptome atlas of the mouse iris at single-cell resolution defines cell types and the genomic response to pupil dilation. Elife 10, (2021).

39. Y. Liu, M. Li, M. Fan, Y. Song, H. Yu, X. Zhi, K. Xiao, S. Lai, J. Zhang, X. Jin, Y. Shang, J. Liang, Z. Huang, Chromodomain Y-like Protein-Mediated Histone Crotonylation Regulates Stress-Induced Depressive Behaviors. Biol Psychiatry 85, 635–649 (2019).

40. M. Fan, Y. Liu, Y. Shang, Y. Xue, J. Liang, Z. Huang, JADE2 Is Essential for Hippocampal Synaptic Plasticity and Cognitive Functions in Mice. Biol Psychiatry 92, 800–814 (2022).

